# Multiplexed profiling of transcriptional regulators in plant cells

**DOI:** 10.1101/2025.05.12.653590

**Authors:** Simon Alamos, Lucas Waldburger, Amanda Dee, Lauren Owens, Rohan Rattan, Shirlyne Ong, Patrick M. Shih

## Abstract

Transcriptional regulators play key roles in plant growth, development, and environmental responses; however, understanding how their regulatory activity is encoded at the protein level has been hindered by a lack of multiplexed large–scale methods to characterize protein libraries *in planta*. Here, we present ENTRAP-seq (Enrichment of Nuclear *Trans*-elements Reporter Assay in Plants), a high-throughput method that introduces protein-coding libraries into plant cells to drive a nuclear magnetic sorting-based reporter, enabling multiplexed measurement of regulatory activity from thousands of protein variants. Using ENTRAP-seq and machine learning, we screened 1,495 plant viruses and identified hundreds of novel putative transcriptional regulatory domains found in structural proteins and enzymes not associated with gene regulation. In addition, we combined ENTRAP-seq with machine-guided design to engineer the activity of a plant transcription factor in a semi-rational fashion. Our findings demonstrate how scalable protein function assays deployed *in planta* will enable the characterization of natural and synthetic coding diversity in plants.

## Introduction

The accelerating pace of plant genomics has facilitated structure- and homology-based comparative approaches to infer the function of proteins regulating important traits; however, the functional annotation of this genetic variation has lagged behind. It remains very difficult if not impossible to predict gene function from sequence alone. Meeting this challenge is a stepping stone towards rationally designing more resilient crop varieties via gene editing^1^. This gap is particularly acute for transcriptional regulators, a class of proteins involved in virtually all aspects of plant biology^2^. Sequence-to-function prediction of transcriptional regulators is hindered by the high disorder and poor conservation of their regulatory domains^3–5^, which limits the power of comparative genomic algorithms to decipher how regulatory activity is encoded at the protein level. Moreover, mutations in regulatory domains are well tolerated or have subtle phenotypic effects, which has conspired against the use of classic random mutagenesis screens^6^. Directed mutagenesis studies are often guided by structural data and sequence conservation, but designing effective mutagenesis experiments becomes challenging when these constraints are absent and the search space is too large.

In recent years, techniques like STARR-seq^7^, DAP-seq^8^, and TARGET ^9^ have enabled mapping the functional requirements of DNA sequences bound and regulated by plant transcription factors (TFs). However, the large-scale *in planta* characterization of the regulatory activity of multiple TFs simultaneously remains challenging and labor-intensive ^10^. Given the limitations of traditional methods, the high-throughput study of TF regulatory domains has focused on exploring their protein sequence space using massively parallel reporter assays (MPRAs) in cultures of animal or microbial cells^11^. In this approach, TF libraries are transformed into cultured cells where they drive the expression of a reporter gene. Cells are then sorted according to the level of reporter expression and next-generation sequencing is used to determine the abundance of each transgene in each sorted bin, a proxy for their activity^12^. These assays empirically characterize transcriptional regulators at a large scale and can yield the breadth of data necessary to train predictive models of the location and function of TF regulatory domains^13–15^. Such models provide a way to perform inference on protein sequences that are too costly or time-consuming to characterize experimentally^16^.

Over the last decade, reporter-based MPRAs of protein libraries in microbes such as *S. cerevisiae* (yeast) and animal cell cultures have opened whole new avenues of inquiry and led to a number of predictive models to predict TF activity^13–15,17–23^. Although proteins of plant origin have been studied in a high throughput manner in yeast^15,19^, these assays have so far not been conducted in plant systems. As a result, our view of plant transcriptional regulation in general and of plant TFs in particular is inherently biased by non-plant model organisms that diverged from plants more than 1 billion years ago^24^. These assumptions are inextricably embedded in current models to predict sequence features dictating the activity of plant TFs^15^. More broadly, MPRAs can be used to study other classes of *trans*-acting and protein-coding genetic elements beyond TFs such as immune effectors or metabolic enzymes whose activity in plants depends on other plant-specific proteins^25^. There is therefore a need for a high-throughput method to measure protein-coding gene libraries expressed in a plant host.

Here, we introduce ENTRAP-seq (<u>E</u>nrichment of <u>N</u>uclear *<u>T</u>rans*-elements <u>R</u>eporter <u>A</u>ssay in <u>P</u>lants), an MPRA based on magnetic sorting of isolated nuclei and the first MPRA capable of measuring *trans*-acting coding variants in plant cells. We deployed this method in *N. benthamiana* leaves expressing gene libraries delivered by *Agrobacterium tumefaciens*, allowing us to simultaneously measure the transactivation capacity of at least 2,000 protein variants *in planta*. We combined this technology with a custom machine learning pipeline to predict and experimentally validate activation domains (ADs) from proteins across 1,495 plant viral genomes. We identified hundreds of peptides of viral origin capable of activating transcription when recruited to a reporter. These potential ADs exhibit positional conservation and tend to occur in structural proteins and enzymes not known to be involved in host gene regulation, such as capsid proteins and RNA polymerases. Moreover, we demonstrate that many of these viral ADs can be used as novel DNA parts for plant synthetic biology, opening the door to large-scale mining of parts optimized for plant engineering. Comparing the activity of plant TF sequences between yeast and *N. benthamiana* revealed pervasive discrepancies between these two systems, challenging current assumptions about AD conservation and highlighting the need for *in planta* assays. Finally, we show that ENTRAP-seq can be used to map, dissect, and engineer the ADs of plant TFs. We used machine-based design to guide semi-rational mutagenesis of the central transcriptional activator in photoperiodic flowering, CONSTANS, identifying sequence features critical for its function and enabling the generation of a new allelic series that includes repressors. ENTRAP-seq bridges a critical methodological gap in plant biology by enabling the multiplexed characterization of natural and synthetic coding variation in plant cells.

## Results

### Optimizing a magnetic nucleus sorting tag as a quantitative reporter assay

To measure the activity of multiple protein-coding elements in parallel, we sought to develop a MPRA based on separating nuclei by the level of expression of a reporter transgene (Fig. 1A). By placing this reporter under the control of a protein-coding library of regulators (i.e., *trans*-regulatory elements), we aimed to sort nuclei into two fractions: a pulldown fraction enriched with regulators that activate the reporter, and a flow-through fraction enriched with repressors and/or inactive elements. Next-generation sequencing is used to quantify the relative enrichment of each *trans-*element in each fraction, which is proportional to their activity (i.e., activators or repressors) and strength (Fig. 1A). As the expression platform, we used *Agrobacterium*-mediated transient expression in *N. benthamiana*, also known as agroinfiltration (Fig. 1A). Crucially, it was previously shown that transgene expression in this system follows Poisson-like statistics^26–28^. This makes it possible to minimize the number of cells expressing more than one *trans*-element from the library by infiltrating *Agrobacterium* cultures at a very low density. Single transgene expression can mitigate unwanted interactions that may occur when different *trans*-elements are expressed in the same cell.

**Figure 1:**
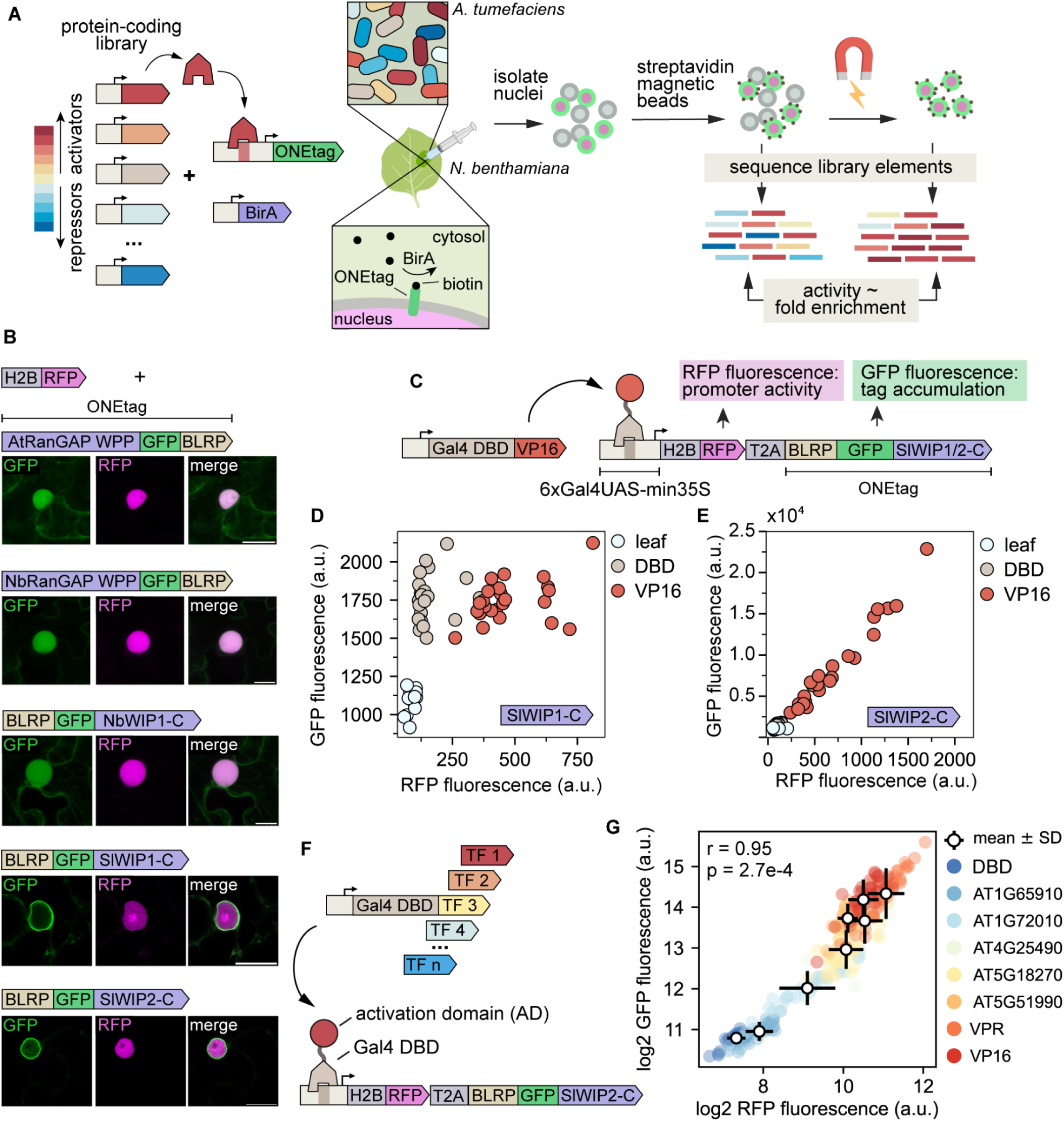
Design and optimization of an assay to measure the activity of *trans*-acting elements in parallel in plant cells. **(A)** Overview of the ENTRAP-seq assay. A library of n different *trans*-acting elements such as transcription factors (TFs) of varying strengths is cloned in binary vectors for *Agrobacterium*-mediated transformation. These *trans*-elements regulate a promoter driving the expression of an <u>o</u>uter <u>n</u>uclear <u>e</u>nvelope tag (ONEtag) containing a biotin acceptor peptide. A transgene for the constitutive expression of the biotin ligase BirA is also included to biotinylate the ONEtag. The library as well as the ONEtag reporter and BirA are transformed into *Agrobacterium* and mixed cultures are infiltrated into leaves of *N. benthamiana*. The ONEtag is then biotinylated by constitutively expressed BirA using endogenous biotin. Next, nuclei are isolated from infiltrated leaves and incubated with magnetic streptavidin-coated beads, allowing magnetic sorting. DNA is extracted from the unsorted and sorted nuclei samples and the abundance of each of the *trans*-elements in the library is determined by sequencing the T-DNA. The enrichment of a given *trans*-element in the sorted vs unsorted population correlates with its activation strength. **(B)** Live cell fluorescence microscopy images of *N. benthamiana* epidermis cells expressing a nuclear marker (H2B-RFP) and different ONEtag candidate versions fused to GFP. Bar = 20 um. Shown on top of each set of images is a schematic of the ONEtag variant tested. **(C)** Schematic of the construct used in (D) and (E) to compare promoter activity (RFP fluorescence) with ONEtag accumulation (GFP fluorescence). **(D)** Scatter plot showing RFP (x axis) and GFP (y axis) fluorescence of leaf punches expressing the construct in (C) containing the SlWIP1-C ONEtag combined with a second strain carrying Gal4DBD or Gal4-VP16. **(E)** Same as (D) for a ONEtag containing SlWIP2-C. **(F)** Schematic of the experiment to test the sensitivity of the SlWIP2-C ONE tag. Gal4DBD fusions to effector domains of Arabidopsis TFs of varying strengths were used to drive the expression of the reporter. **(G)** Scatter plot of the GFP (x axis) and RFP (y axis) fluorescence expressing the constructs in (F). In (D), (E), and (G) N = 24 (3 plants with 8 leaf punches per plant).

To link reporter expression to nucleus sorting, we used a chimeric tag composed of a fusion of a biotin acceptor peptide to GFP and an <u>o</u>uter <u>n</u>uclear <u>e</u>nvelope targeting domain (hereafter ONEtag) which can be biotinylated by a coexpressed BirA transgene using endogenous biotin. Nuclei extracts incubated with streptavidin magnetic beads can then be sorted using magnets. Similar approaches have been used to purify nuclei from specific cell types in transgenic plants^29^, but there are no reports of magnetic sorting of transiently transformed *N. benthamiana* tissues. We hypothesized that transiently transformed nuclei could be sorted into two fractions: a pulldown fraction enriched in T-DNAs encoding activators, and a flow-through fraction depleted of strong activators and enriched in T-DNAs coding for repressors and neutral elements (Fig. 1A). Importantly, for this approach to be quantitative, magnetic nucleus sorting must be sensitive enough to detect quantitative differences in ONEtag expression levels among cells, rather than merely distinguishing between nuclei that do and do not express the ONEtag, as has been previously done to isolate cell type-specific nuclei^30^. To optimize magnetic nucleus sorting as a quantitative readout of transcriptional activity in agroinfiltrated *N. benthamiana*, we pursued two design goals. First, the transiently expressed ONEtag should localize to the nuclear envelope, as determined by the GFP subcellular localization. Second, as for any sensitive quantitative reporter, the level of ONEtag accumulation must correlate with the transcriptional activity of the promoter driving its expression.

As the first candidate ONEtag, we tested the AtRanGAP WPP domain, previously used to label nuclei in specific tissues in *Arabidopsis*. To confirm ONEtag subcellular localization, we co-expressed a nuclear marker (pUBQ10::H2B:mScarlet, hereafter H2B-RFP). When transiently expressed in *N. benthamiana*, the WPP tag did not localize to the nuclear envelope, as indicated by its failure to encircle the H2B-RFP signal (Fig. 1B). A similar pattern was observed for the WPP domain of the closest *N. benthamiana* RanGAP homolog. Since the C-terminus of WPP proteins has also been used to target the nuclear envelope, we tested the C-terminal half of the *N. benthamiana* homolog of AtWIP1, which also failed to localize to the nuclear envelope. We next screened seven domains known to localize to the outer nuclear envelope in their native contexts, including the C-termini of two tomato WIP proteins. Several candidates showed correct localization in *N. benthamiana* (Fig. S1A), and we selected the C-terminal halves of WIP1 and WIP2 (SIWIP1/2-C) for further characterization (Fig. 1B).

We next focused on the second design criterion, that is, ensuring that the degree of ONEtag protein accumulation correlates with the transcriptional activity of the promoter driving its expression. To measure promoter activity independent of ONEtag protein abundance, we used an RFP-T2A-ONEtag fusion (Fig. 1C). RFP fluorescence intensity served as a proxy for promoter activity while GFP fluorescence intensity provided a metric for ONEtag accumulation. To drive the RFP-T2A-ONEtag reporter, we placed this transgene under the control of a minimal 35S promoter with 6 upstream Gal4 DNA binding sites (UAS-min35S-RFP-T2A-ONEtag). To activate expression of this reporter, we coinfiltrated it in *N. benthamiana* with previously generated strains containing the Gal4 DNA binding domain alone (Gal4DBD) or fused to the strong VP16 activator (Gal4-VP16)^10^. As expected, the level of RFP fluorescence was higher in leaves coexpressing the SlWIP1-C ONEtag and Gal4DBD-VP16 compared to Gal4DBD (Fig. 1D). However, GFP fluorescence was comparable between Gal4DBD-VP16 and Gal4DBD, suggesting that the increase in transcriptional activity driven by the VP16 fusion was buffered postranscriptionally, preventing an increase in SlWIP1-C ONEtag protein accumulation (Fig. 1D). Cell microscopy confirmed that the ONEtag and RFP proteins are efficiently separated after being produced from the same transcripts (Fig. S1B). While removal of BLRP did not relieve the buffering effect, removing SlWIP1-C was sufficient for GFP fluorescence to increase in response to VP16, similar to RFP (Fig. S1C,D). We therefore replaced SlWIP1-C with SlWIP2-C. For this ONEtag, GFP and RFP accumulated linearly in response to VP16 showing that the accumulation of this tag closely correlates with the activity of the promoter driving it (Fig. 1E). Removing RFP-T2A from the constructs made accumulation of SlWIP1-C ONEtag slightly responsive to Gal4-VP16 but to a far lesser degree than SlWIP2-C (Fig. S1D) To confirm that the accumulation of the SlWIP2-C ONEtag tracks with promoter activity across a large dynamic range, we coexpressed it with 8 previously generated Gal4DBD fusions spanning a wide range of transcriptional strengths^10^. Across combinations, RFP and GFP fluorescence correlated well (R^2^ = 0.91, Fig.1F,G Fig.S2, Table S1). Hence, accumulation of the SlWIP2-C ONEtag is highly sensitive and quantitatively reflects the transcriptional activity of the promoter driving its expression.

### Small-scale library validation of ENTRAP-seq

Having optimized the purification tag, we next sought to validate that our approach can be used to quantitatively measure the activity of *trans*-acting elements. For enrichment in the nucleus pulldown fraction to serve as a quantitative metric of activity, the probability of pulling down a given nucleus should be correlated with its level of ONEtag expression. For this pulldown bias to yield a quantitative metric of reporter expression, the correlation between promoter activity and ONEtag abundance that we observed at the tissue level should also apply in individual nuclei. The ultimate goal of ENTRAP-seq is to quantitatively measure the level of expression of a reporter. Hence, we sought to benchmark ENTRAP-seq by comparing its output (i.e., TF enrichment in the pulldown based on next-generation sequencing) against that of a well-established fluorescence reporter assay.

To demonstrate that ENTRAP-seq meets these criteria, we characterized a control set of 26 previously generated synthetic TFs consisting of 53 amino acid (aa) fragments –hereafter referred to as ‘tiles’– fused to Gal4DBD (Table S2)^19^ and spanning a wide range of activity, from repressors to strong activators^19^. We cloned these Gal4DBD fusion tiles under the constitutive pMas promoter in an *Agrobacterium* T-DNA expression plasmid containing the SlWIP2-C RFP-T2A-ONEtag construct under the control of UAS-min35S and a constitutive BirA transgene (Fig. 2A). All three cassettes were placed in the same T-DNA to ensure that biotinylated ONEtag expression occurs exclusively in cells that express a synthetic TF. We used this backbone (pSA676v2) for all subsequent ENTRAP-seq experiments and fluorescence reporter assays involving Gal4DBD fusions. We previously determined that a low *Agrobacterium* OD (optical density at 600 nm) of approximately 0.0025 is optimal for expressing libraries of mixed *trans*-elements in *N. benthamiana* as it maximizes the number of transformed cells while minimizing the fraction of these cells that express more than one transgene^26^. At this OD, approximately 10% of transformable leaf cells express a transgene and about 90% of these cells express a single one. For the following nucleus pulldown experiments, we therefore infiltrated the 26 control strains at a total mix OD of 0.0025 (Fig. 2B). To purify and sort nuclei, we developed a protocol based on pipelines optimized for other plant species, with modifications (Materials and Methods)^30,31^.

**Figure 2:**
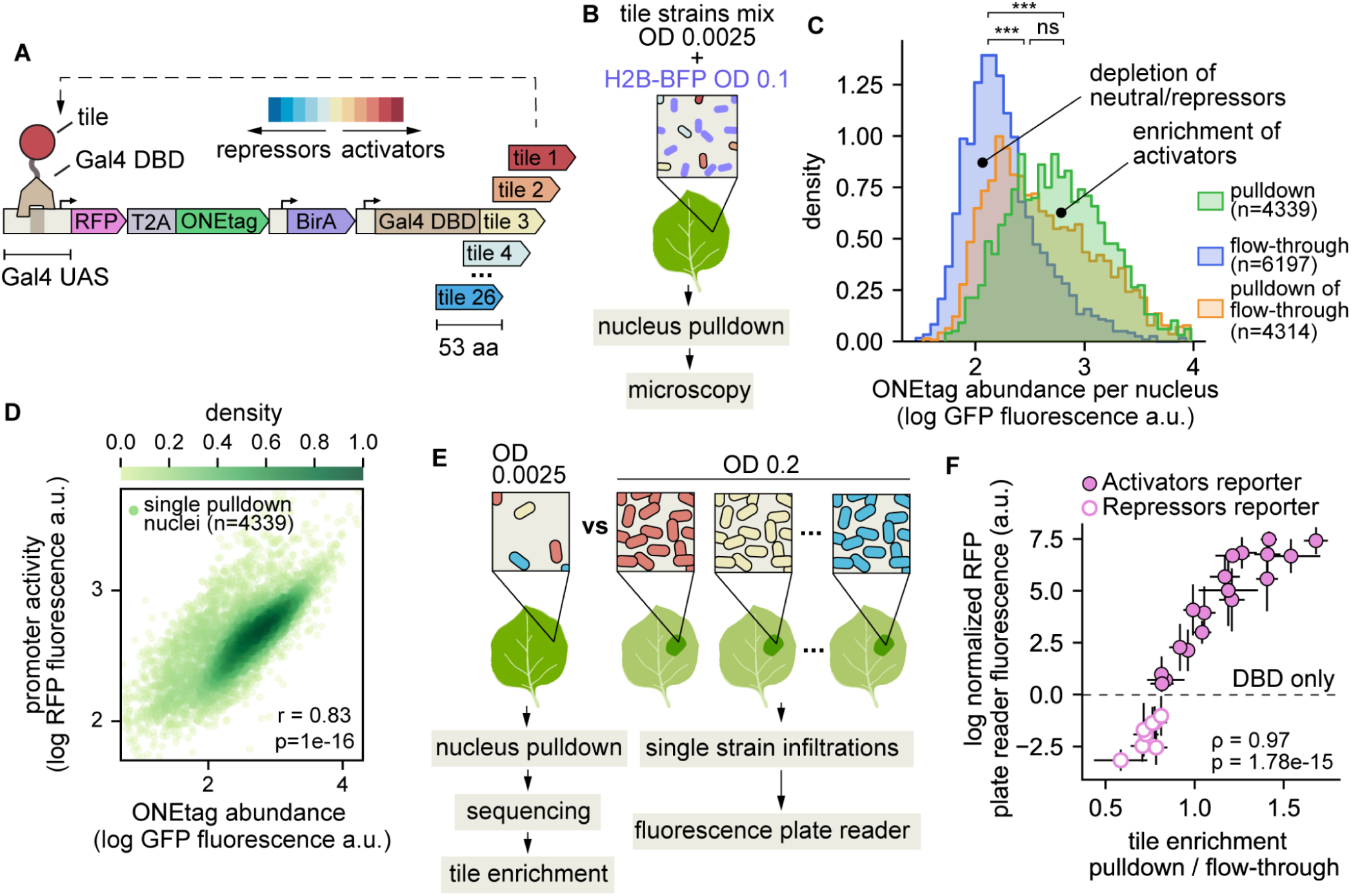
Validation of ENTRAP-seq for quantitative measurements of reporter expression. **(A)** T-DNA constructs used to validate ENTRAP-seq. A synthetic promoter containing 6x Gal4 UAS sites followed by the minimal 35S promoter is used to drive the ONEtag fused to RFP by a self-cleaving T2A peptide. Downstream of the ONEtag, the plasmid contains a constitutive BirA expression cassette and a constitutively-expressed fusion between the Gal4 DBD and variable 53 aa tiles of various strengths. **(B)** Infiltration setup for nucleus pulldown and microscopy experiments shown in (C) and (D). Strains carrying the 26 constructs from (A) are grown separately and mixed in equal OD ratios prior to leaf infiltration at a final total OD of 0.0025. This mix is combined with a strain carrying a constitutive H2B-BFP nuclear marker infiltrated at OD 0.1. This tissue is then subjected to magnetic nucleus pulldown and the BFP signal is used to identify the nuclei of transformable leaf cells. **(C)** Distribution of ONEtag expression levels in nuclei from different purification fractions. Histograms are normalized to have the same area. Nuclei with low GFP levels are enriched in the flow-through and there is a progressive enrichment in the pulldown with increasing GFP expression. *** p<0.001, ns distributions not significantly different using Mann–Whitney U test. **(D)** Scatterplot of RFP versus GFP fluorescence in single nuclei in the pulldown fraction from (C). **(E)** Schematic of the experiment to compare ENTRAP-seq enrichment with fluorescence reporter assays. Left: the control set of 26 tiles from (A) was infiltrated at a low OD for nucleus pulldown and sequencing of the flow-through and pulldown fractions. Right: each of the strains in (A) is infiltrated individually at an OD of 0.2. The activity of each tile is determined by measuring the RFP fluorescence of infiltrated leaves in a plate reader. **(F)** X-axis: transcriptional activity of each tile based on ENTRAP-seq (mean ± SD enrichment of each tile T-DNA in the nuclei pulldown fraction compared to the flow-through sample). Y-axis: Tile transcriptional activity measured individually using a fluorescence plate reader (mean ± SD RFP fluorescence). N for nuclei pulldown and sequencing = 4 biological replicates. N for plate reader assays = 24 (3 plants with 8 leaf punches per plant). ⍴ = Spearman correlation coefficient, r = Pearson’s correlation coefficient.

To fluorescently label transformable nuclei in the different purification fractions, we infiltrated the tile mix with an H2B-BFP strain at OD 0.1, which ensures homogeneous and strong labeling across cells (Fig. 2B). Using the BFP nuclear marker, we counted nuclei in the pulldown and the flow-through fractions and calculated their level of ONEtag expression by measuring GFP fluorescence (Materials and Methods). Using this approach we counted approximately 300,000 nuclei in the pulldown fraction starting from 10g of fresh tissue (about 20 fully infiltrated leaves) (Fig. S3A). The flowthrough fraction contained approximately 10 million nuclei (Fig. S3A). These numbers are within a factor of 2 from an estimate of the total number of transformed cells in the sample (see estimation in Fig. S3B). As shown in Figure 2C, nuclei in the pulldown had a higher level of ONEtag expression than those in the flow-through, while a second pulldown showed an intermediate distribution (Fig. 2C). These distributions show a large degree of overlap such that the relative abundance in the pulldown with respect to the flow-through increases with ONEtag abundance (Fig. 2C). This feature is critical for the pulldown-to-flowthrough ratio to serve as a continuous metric of transcriptional strength. We obtained qualitatively similar results in a biological replicate of this experiment (Fig. S3C). Comparing mCherry and GFP fluorescence at the single nucleus level revealed strong correlation between reporter activity and ONEtag accumulation (Fig. 2D, Pearson’s r = 0.83), further validating the premise of our method.

Next, we sought to directly demonstrate that ENTRAP-seq can be used to quantitatively measure reporter activity. To this end, we asked whether the degree of pulldown enrichment of each of the 26 control tiles correlates with their transcriptional strength measured using an established reporter assay. We infiltrated each construct individually and used a fluorescence plate reader to measure the tissue RFP and GFP fluorescence normalized to a constitutive BFP strain. This ratio was then expressed as a function of the no-tile Gal4DBD control. In contrast to the mixed infiltrations for nucleus pulldown, single tile infiltrations for fluorescence measurements were done at OD 0.2 to ensure a strong fluorescence signal (Fig. 2E, Materials and Methods: Fluorescence plate reader reporter assays). The basal fluorescence of the min35S promoter is close to leaf autofluorescence, hindering the detection of transcriptional repression in fluorescence reporter assays (Fig. S4). Hence, in the case of repressor tile strains we coinfiltrated them with a third construct containing a synthetic promoter with embedded Gal4 binding sites, which has a ∼4x higher basal expression level than min35S^32^ (Fig. S4). To measure the ENTRAP-seq enrichment of these tiles we combined the corresponding 26 *Agrobacterium* cultures in equal OD ratios and infiltrated the mix at a final OD of 0.0025. Nuclei purified from these leaves were subjected to magnetic sorting and total DNA was extracted from the pulldown and flow-through fractions. Amplicon sequencing of the T-DNA tile region was used to determine the relative abundance of each tile in each fraction in order to obtain their level of enrichment. Tile enrichment correlated very well with promoter activity as measured by leaf RFP fluorescence (Spearman ⍴ = 0.97), indicating that ENTRAP-seq can accurately and quantitatively measure TF regulatory strength (Fig. 2F). As expected, leaf GFP fluorescence also correlated well with tile enrichment (Fig. S3D, Spearman ⍴ = 0.97) and with RFP fluorescence (Fig. S3E, Pearson’s r = 0.96).

Overall, these results demonstrate that by linking reporter activity to relative pulldown enrichment, ENTRAP-seq provides a method to quantify *trans*-element activity in plant cells.

### Accuracy and reproducibility in large libraries of viral protein tiles

We next sought to leverage ENTRAP-seq to discover novel potential transcriptional regulators. Plant pathogen attack is usually linked to widespread transcriptomic changes in the host through the action of pathogen-encoded transcriptional regulators^33^. There are well-known examples of such regulators delivered by pathogenic bacteria into plant cells, but how plant viruses manipulate host transcription is much less understood^34^. In the case of DNA viruses, transcriptional regulation of the viral genome is part of the pathogenic cycle. Plant viruses have very small genomes (typically in the order of 10-15 Kbp, Fig. S5) that encode just a few proteins (normally about 4, Fig. S5), none of which resemble known transcription factors in terms of sequence homology and presence of known TF domains^35^. Notably, many of the most widely used genetic parts in plant and mammalian genetic engineering – e.g., the VP16 AD, the 35S promoter, and the SV40 NLS – originate from plant and animal viruses. More broadly, a better molecular understanding of the viral transcriptional arsenal could help devise new crop protection strategies.

To identify potential viral transcriptional regulators, we searched the plant viral proteome for putative ADs. Throughout this study, we adopted a strictly functional definition of ADs, namely, short peptides which are sufficient for strongly activating transcription of a promoter when recruited via a heterologous DBD. Importantly, the fact that a peptide of viral origin fits this definition does not necessarily imply that its protein of origin is involved in gene regulation nor that this peptide is required for regulating transcription within its protein context. Indeed, many potent ADs are found in non-nuclear proteins that lack known gene regulatory functions, such as metabolic enzymes in mitochondria and chloroplasts^19^. A peptide may adopt a different structural conformation when isolated and fused to a DBD compared to its native protein context. As a result, tiles derived from TFs can sometimes behave as ADs in reporter assays even though they seem dispensable for the transactivation function of the TF itself. This functional definition is meant to emphasize that AD identification in reporter recruitment assays is but a first step toward fully characterizing the regulatory function of a protein.

As of the time of writing, the NCBI database contained 1,495 plant viral genomes and 5,607 proteins. To maximize sequence diversity, we first clustered the protein sequences at 90% sequence identity, yielding 4,537 representative nonredundant proteins. Each sequence was then tiled using 53 aa windows and 10 bp overlaps (Fig. 3A). To estimate the likelihood of each tile being a putative AD, we used PADDLE, a deep convolutional neural network that predicts acidic ADs^14^. The convolutional layer enables PADDLE to identify sequence-to-function relationships, especially within disordered sequences, where traditional structure- or homology-based comparative genomic algorithms fail. Although the model was trained on data from a yeast-based screen using yeast protein sequences, acidic ADs are evolutionarily-conserved and we previously demonstrated that PADDLE can identify ADs from plants^19^. Only tiles with the highest predicted z-scores were selected for experimental testing (Fig. 3A). Finally, we further optimized protein diversity by selecting sequences that maximized the Hamming distance between binary protein encodings (Fig. 3A).

**Figure 3:**
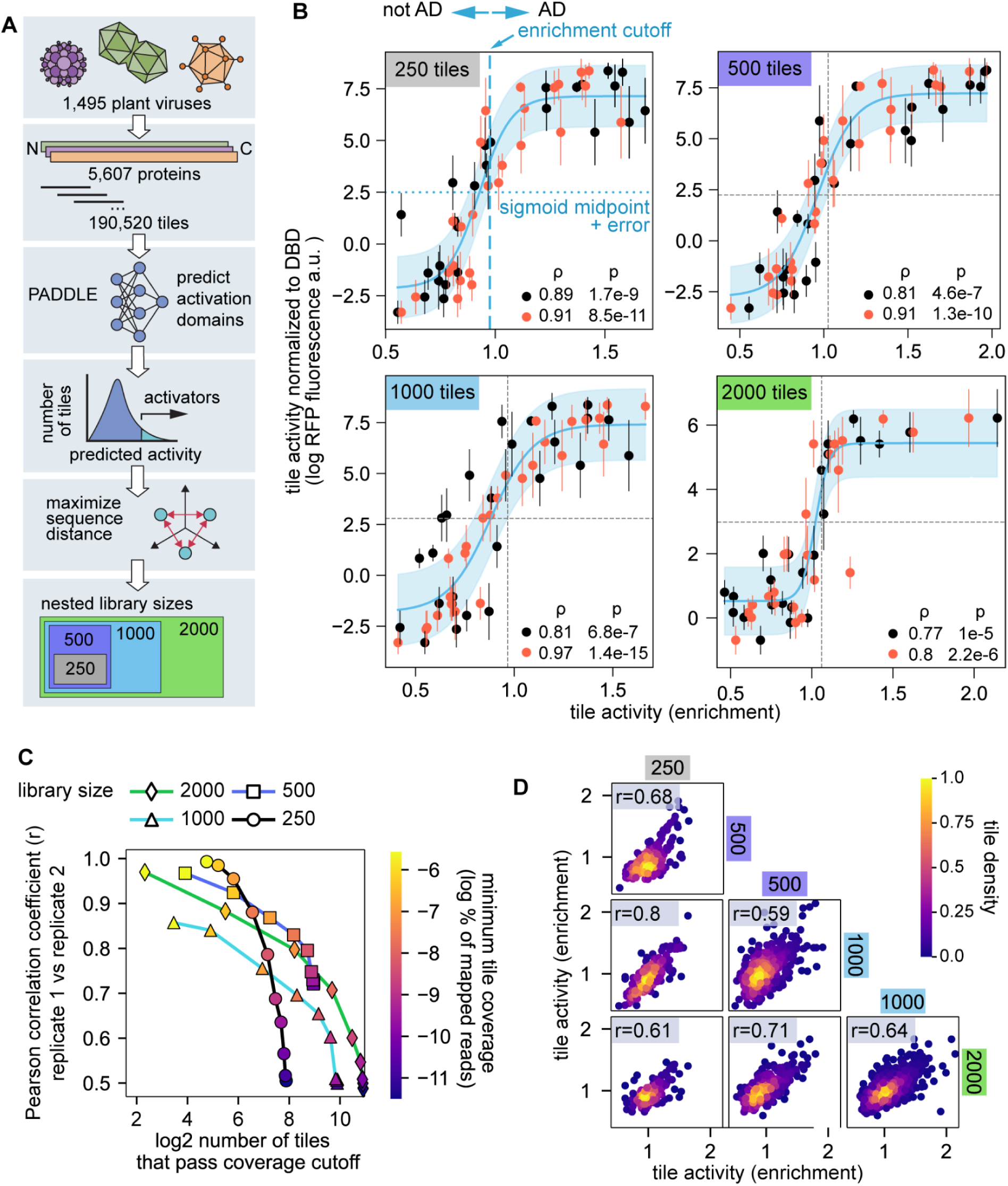
Generation of nested libraries of putative viral activation domains to test assay scale and reproducibility. **(A)** Schematic of the computational pipeline to generate libraries of putative transcriptional activation domains from plant viruses. From the top: protein ORF sequences from genomes of all plant viruses in NCBI were deduplicated and clustered using 90% sequence identity then fragmented into 53 aa tiles with 10 bp overlap. PADDLE was used to predict the transactivation strength of these tiles. The top 2,000 tiles with the highest predicted activity were selected (PADDLE z-score > 4.9). Protein sequences were binary encoded, and sequence diversity maximization algorithm was applied to iteratively select sequences that maximized the average Hamming distance from those already included in the subset. This approach enriched for sequence-level diversity in the selected proteins to generate nested libraries of 250, 500, 1K, and 2K tiles. **(B)** Tile activity as measured by ENTRAP-seq (x axis, sequencing-based enrichment) plotted against the mean ± 1 s.d. tile activity measured individually using a fluorescence reporter assay (y-axis) for two biological replicates (orange and black circles) of a library of 250, 500, 1K, and 2K viral tiles. Blue lines show a sigmoidal fit to the data with the shaded blue area indicating ± 1 s.d. of residuals as an empirical confidence bound. Horizontal dashed lines indicate the middle point of the sigmoid fits. Vertical dashed lines indicate the enrichment value corresponding to the sigmoid midpoint + 1 s.d. This value was taken as the lower bound of enrichment to classify tiles as activators. **(C)** Reproducibility between biological replicates (Pearson correlation in enrichment, y axis) as a function of the absolute number of tiles that pass a threshold number of mapped reads (x axis). The set of tiles counted on the x axis was generated by imposing coverage cutoffs such that only tiles with more counts than an arbitrary and variable threshold (color bar) were considered to calculate the Pearson correlation. **(D)** Reproducibility across different libraries. The activity of each tile was first averaged across both biological replicates of each library of a given size. Scatter plots show the activity of tiles that are shared between libraries of different sizes. The Pearson correlation (r) between libraries is shown.

Although our nuclei enrichment method performed well on the control set of 26 tiles (Fig. 2F), the scalability of ENTRAP-seq to larger libraries was not immediately clear. To address this question, we created 4 nested libraries of 250, 500, 1K, and 2K viral tiles (Fig. 3A). All except for the 2K library also contained the 26 control tiles for which we had previously obtained single infiltration fluorescence data (Fig. 2F). For the 2K library, a new set of 26 viral tiles was measured individually in a fluorescence plate reader. These internal controls allowed us to validate the output of the assay by comparing the activity of these tiles in terms of reporter fluorescence and tile enrichment (as in Fig. 2F). Across all libraries, we found that the activity of a given control tile as measured individually in a fluorescence plate reader strongly correlates with its enrichment following magnetic pulldown for two biological replicates of each library (Spearman ⍴ = 0.72-0.97) (Fig. 3B-E). These validation experiments also allowed us to obtain a calibration value to classify tiles as activators based on their sequencing-based enrichment. To this end, we fitted the validation data to a sigmoidal curve and calculated the enrichment value corresponding to the midpoint of this sigmoid plus its estimation error (Fig. 3B-E). This yielded an enrichment value of approximately 1.1, which was used as a cutoff to classify viral tiles as activators in subsequent analyses (Fig. 3B-E).

We found no consistent correlation between tile coverage in the library and tile enrichment ratio (Fig. S6). When the enrichment of all tiles was considered, we found that biological replicates correlated well (Pearson r = 0.5-0.55) and this correlation increased substantially to r = 0.7-0.8 by applying even modest library coverage cutoffs (Fig. 3C). Because of the nested nature of the libraries, it was also possible to compare tile enrichment across libraries. This revealed a high degree of reproducibility even across libraries (Pearson’s r = 0.59-0.8 without applying coverage cutoffs) (Fig. 3D). Thus, our assay is accurate and reproducible in libraries of up to 2K tiles.

### Identification of putative activation domains in plant viral proteins

Our pipeline for discovering potential viral ADs combining machine learning predictions with high-throughput measurements yielded 1,105 tiles that passed the activator enrichment cutoff, confirming that PADDLE can predict acidic ADs in plants^19^ (Fig. 4A, Fig. S7-10). These tiles are found throughout the phylogenetic tree of plant viruses (Fig. 4A). Two large families of plant ssDNA and dsDNA viruses (*Geminiviridae* and *Caulimoviridae*, respectively) as well as multiple +ssRNA families (especially *Potyviridae* and *Betaflexiviridae*) were particularly rich in acidic ADs (Fig. 4A). We did not find any ADs in only 4 relatively small families: *Amalgaviridae*, *Aspiviridae*, *Mitoviridae*, and *Nanoviridae* (-ssRNA, -ssRNA, +ssRNA, and ssDNA, respectively). This contrasts with recent global surveys of human viruses where ADs were found almost exclusively in a few families of dsDNA viruses^22,36^. Although our screen did not encompass all viral genomes, it identified approximately twice as many tiles as the more comprehensive studies of human viral activation domains, which reported ∼600 ADs. This highlights the utility of using a predictive model like PADDLE to bias the AD search when it is not technically feasible to explore the whole sequence space. In general, our search algorithm based on PADDLE performed remarkably well in terms of predicting ADs; ∼50% of the tested tiles qualify as activators based on our calibration thresholds, although there was substantial variation in predictive accuracy across taxa (Fig. 4A, Fig. S7-10).

**Figure 4:**
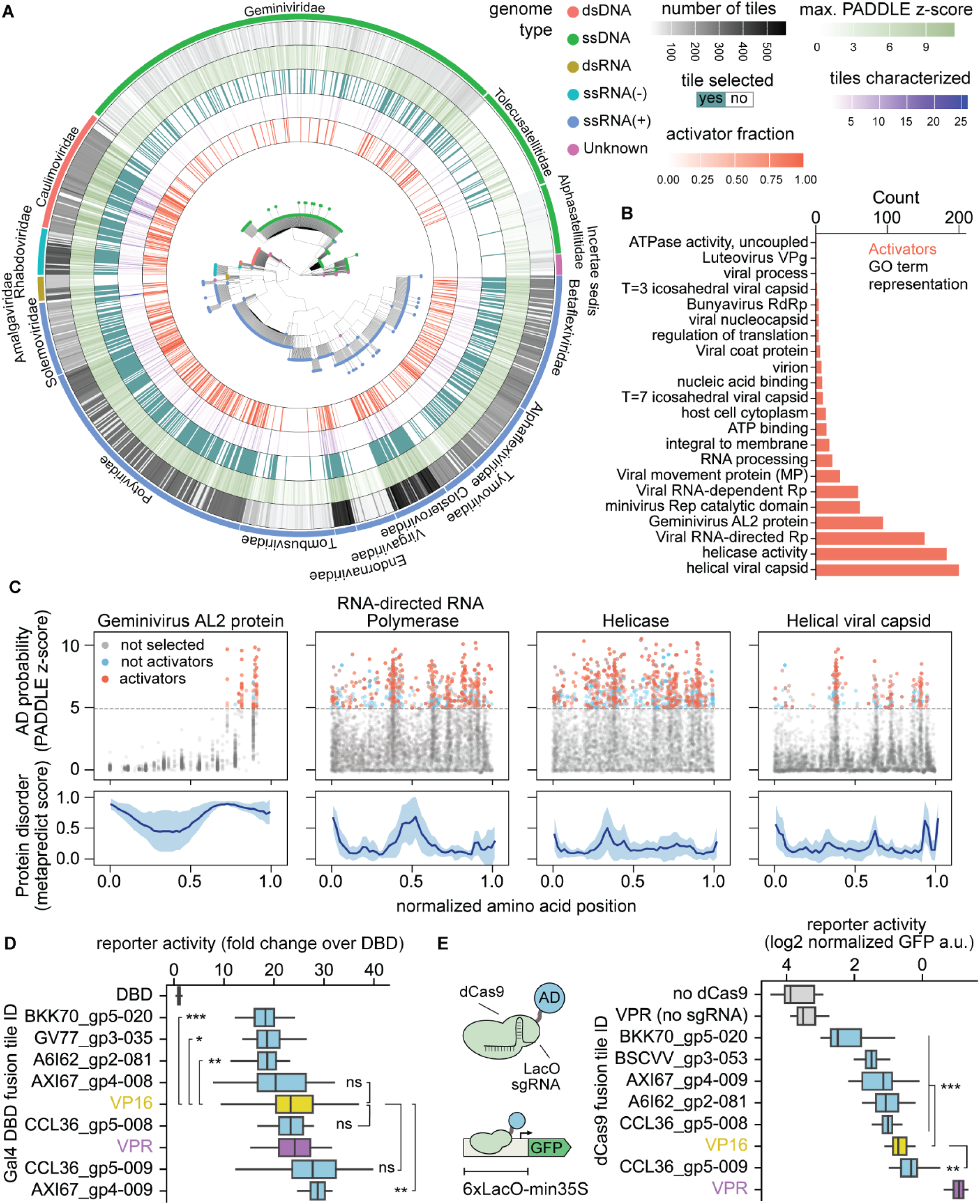
Conserved and potent transcriptional activation domains are found in diverse proteins across plant virus taxa. **(A)** Phylogenetic tree of plant viruses showing the number of tiles selected from each viral genome (gray), the maximum PADDLE predicted z-score (light green), whether a tile from this viral genome was selected for characterization (dark green), the number of tiles that were experimentally measured (purple), and the proportion of measured tiles that were activators (orange). **(B)** Gene ontology (GO) enrichment analysis of tiles measured as activators across all libraries. **(C)** Detailed view of tiles from the proteins most enriched in activator tiles in (B). Top: AD prediction (PADDLE z-score) of each tile along the length of all proteins belonging to each given class. The amino acid position of each tile was normalized by dividing by the length of the protein it originates from. The horizontal dashed line shows the z-score cutoff used to select tiles for testing. Blue dots show tiles that were experimentally tested but did not pass our activator enrichment cutoff. Orange dots show tiles that were tested and had an enrichment higher than the cutoff. Bottom: average ± 1 s.d. protein disorder as predicted by Metapredict. **(D)** Selected strong activator viral tiles tested individually in *N. benthamiana* using a Gal4DBD recruitment fluorescence reporter assay. Shown is the GFP fluorescence normalized to the Gal4DBD alone (Materials and Methods: Fluorescence plate reader reporter assays). **(E)** A set of tiles tested in (D) were tested by recruiting them to a synthetic reporter driving GFP using dCas9 fusions. Shown is the normalized leaf GFP fluorescence. In (D) and (E) brackets indicate two-sided Welch’s t-test comparisons against VP16 with a Bonferroni multiple comparison correction. * p < 0.05, ** p < 0.01, *** p < 0.001, ns not significant. In (D) and (E) box plots show the median ± IQR, whiskers correspond to minimum and maximum values excluding outliers.

To better understand the origin of these potential regulators, we performed GO term enrichment analysis. We found activator tiles in proteins from a wide range of functional annotations, although we note that due to the biased nature of our search not all functional classes are represented equally in our libraries (Fig. 4B, Fig. S11). The presence of ADs in structural and catalytic proteins may reflect the multifunctional nature of viral proteins and mirrors the findings of a recent study that tiled human viruses, which found that 90% of putative viral ADs are found in nonregulatory proteins ^22^. Four kinds of plant viral proteins showed a notable enrichment of activator tiles: geminiviral AL2-like proteins, RNA-dependent RNA Polymerases (RdRp), helicases, and helical capsid proteins (Fig. 4B). The AL2 protein, also known as TrAP (transcriptional activator protein), is involved in transactivating viral promoters in *Geminiviridae* and has been shown to regulate genomic promoters by physically interacting with host TFs^37,38^. AL2 lacks a DNA-binding domain but contains a C-terminal acidic region that can transactivate a synthetic reporter when fused to Gal4DBD^39^. Thus, the recovery of strong ADs in AL2-like proteins using ENTRAP-seq validates our framework for discovering new viral-encoded transcriptional regulators. Our high-resolution tiling of these proteins allowed us to locate the position of their putative ADs (Fig. 4C). Consistent with the C-terminus of AL2 being important for transactivation, we found that ADs are consistently located in three positions within a highly disordered region in the C-terminus of diverse AL2 proteins (Fig. 4C, Fig. S12). The positions of RdRp activator tiles are also relatively conserved but unlike AL2 these putative ADs are clustered in multiple distinct regions throughout the protein, most of which are predicted to fold into stable structures (Fig. 4C, Fig. S13). Similar to RdRp, putative activator tiles in viral helicases and helical viral capsid proteins consistently fall in a few structured regions (Fig. 4C, Fig. S12, Fig. S13). In terms of prediction accuracy, the large majority of predicted ADs in AL2 and helical capsid proteins turned out to be true activators while the opposite was true for RdRp and helicases (Fig. 4C, Fig. S12). These data suggest that a dedicated regulatory protein like AL2 might activate transcription in a similar manner as canonical eukaryotic TFs with disordered ADs, while multifunctional viral proteins may repurpose stably folded and conserved catalytic or structural domains to accomplish this task. Alternatively, the peptides from these nonregulatory proteins may not actually function as ADs in their native context but simply have a sequence composition that led us to serendipitously identify them as activators in our screening setup. This large dataset should serve as a comprehensive starting point for future testing of specific hypotheses regarding the role of these putative ADs in their native protein context.

The discovery of these putative viral regulators prompted us to consider their potential as novel plant synthetic biology tools. To find potent and compact activators that could serve as alternatives to existing field standards, we individually cloned 20 tiles that were among the most enriched in at least one library. We then tested them individually using the same Gal4-UAS fluorescence reporter assay described above, side by side with two potent popular ADs used in the field, VP16 and VPR^40^. All tested tiles activated the reporter at least 20-fold over the DBD alone (Fig. S14). We repeated this experiment for the strongest 7 candidates and verified that 4 performed as well or better than VP16 and VPR (Fig. 4D), confirming that tile enrichment quantitatively captures AD strength. To determine if these ADs perform well in different genetic contexts, we fused them to dCas9 and tested their activity upon recruitment to a synthetic promoter driving GFP as a reporter (Fig. 4E)^41^. dCas9-VPR outperformed all other fusions but one tile was stronger than VP16 when fused to dCas9 (Fig. 4E). Since VPR is considerably larger than our viral tiles (553 vs 53 aa), this compact viral AD may be advantageous for vectors with a limited size capacity such as viral vectors ^42,43^.

To characterize potential new repressor domains of viral origin, we followed a similar strategy, selecting 20 tiles that were highly depleted across libraries (Fig. S14). As field standard controls, we included 4 of the most potent previously characterized synthetic EAR repression motifs ^44^. Only 3/20 of these putative repressors had significantly lower reporter expression than Gal4DBD but none surpassed synthetic EAR motifs in potency. The remaining 17/20 tiles were statistically indistinguishable from the Gal4DBD control (Fig. S14). Thus, the dynamic range of ENTRAP-seq spans from transcriptionally neutral tiles to the most potent known activators but it is not large enough to resolve transcriptional repressors. To specifically detect transcriptional repression, it may be necessary to use a reporter with a higher basal expression level in combination with changes to the pulldown protocol itself ^20^.

In addition to minimizing construct size, these viral ADs offer a diverse set of genetic parts to reduce unintended consequences in synthetic programs, including cross-interactions with modules sharing identical parts and gene silencing arising from context-dependent effects.

### Partial conservation of AD function between distantly related eukaryotes

Recently, plant protein sequences have been used in massively parallel reporter assays in yeast to identify peptides that are sufficient for transcriptional activation in this model system^15,19^. Deriving insight about plant ADs from these experiments in a distantly-related host depends on the degree to which *trans*-activation mechanisms are conserved between these two systems. It is well documented that there is conservation of transcriptional activation mechanisms among eukaryotes, including between yeast and plants^10,45^. Decades of insights about plant TFs derived from yeast-based assays and our own results showing that a model trained in yeast successfully predicts plant viral ADs support this idea^19^. On the other hand, plants have highly derived and expanded transcription factor families^46–48^ and plant transcriptional regulation displays many regulatory features not shared by other eukaryotes^19,24,49^. Not surprisingly, yeast-based screens can miss known ADs in well-studied plant TFs ^15^. A survey of tens of ADs found that transactivation potency in yeast does not predict strength in *N. benthamiana*^19,24^ while an earlier study showed that the quantitative activity of mutant variants of a maize TF was largely consistent between yeast and maize^50^. Thus, it is not clear the degree to which yeast and plants have their own taxa-specific activation mechanisms and whether the quantitative activation strength of ADs is consistent between these two models ^19,24^. To systematically study these questions, we took advantage of an existing dataset where thousands of protein tiles from *Arabidopsis* TFs were tested in yeast^15^. We leveraged ENTRAP-seq to test a subset of these tiles in *N. benthamiana* in order to compare their activity across model systems. We randomly selected 250 tiles while keeping the same distribution of yeast transcriptional strengths (referred to as PADI scores^15^) from the original library. We then cloned this library in the same vector used for our viral libraries and performed a nucleus enrichment experiment.

Comparing the rank ordering of activity between yeast and *N. benthamiana* revealed a very limited correlation (Spearman ⍴ = 0.18, p = 0.0075) (Fig. 5A). Indeed, numerous strong yeast activators were depleted from the ENTRAP-seq pulldown and multiple enriched tiles in *N. benthamiana* were not found to be activators in yeast (Fig. 5A). Because yeast-based MPRAs are heavily biased towards identifying acidic ADs, we hypothesized that tiles not found to be activators in yeast might belong to other classes of ADs. To classify tiles as likely plant activators we used an enrichment threshold of 1.2 — the minimum enrichment value for validated activators from the viral libraries. Among the 40 tiles that had an ENTRAP-seq enrichment greater than 1.2 and a PADI score lower than 1 (labeled ‘quadrant 4’ in Fig. 5A) we found two Q-rich sequences that were not classified as activators in yeast. The *trans*-activation role of Q-rich stretches is conserved in eukaryotes^51^. The first of these polyQ ADs is located in the middle region (MR) of ARF19 (Fig. 5B). The MR is known to be important for *trans*-activation in activator ARFs^52^ but the yeast assay mapped the ADs of ARF19 and other activator ARFs to regions flanking the MR^15^. The second polyQ stretch belongs in the C terminus of the AGAMOUS-LIKE gene ALG65, a region that is thought to contain the AD in this family of TFs (Fig. 5C)^53^.

**Figure 5:**
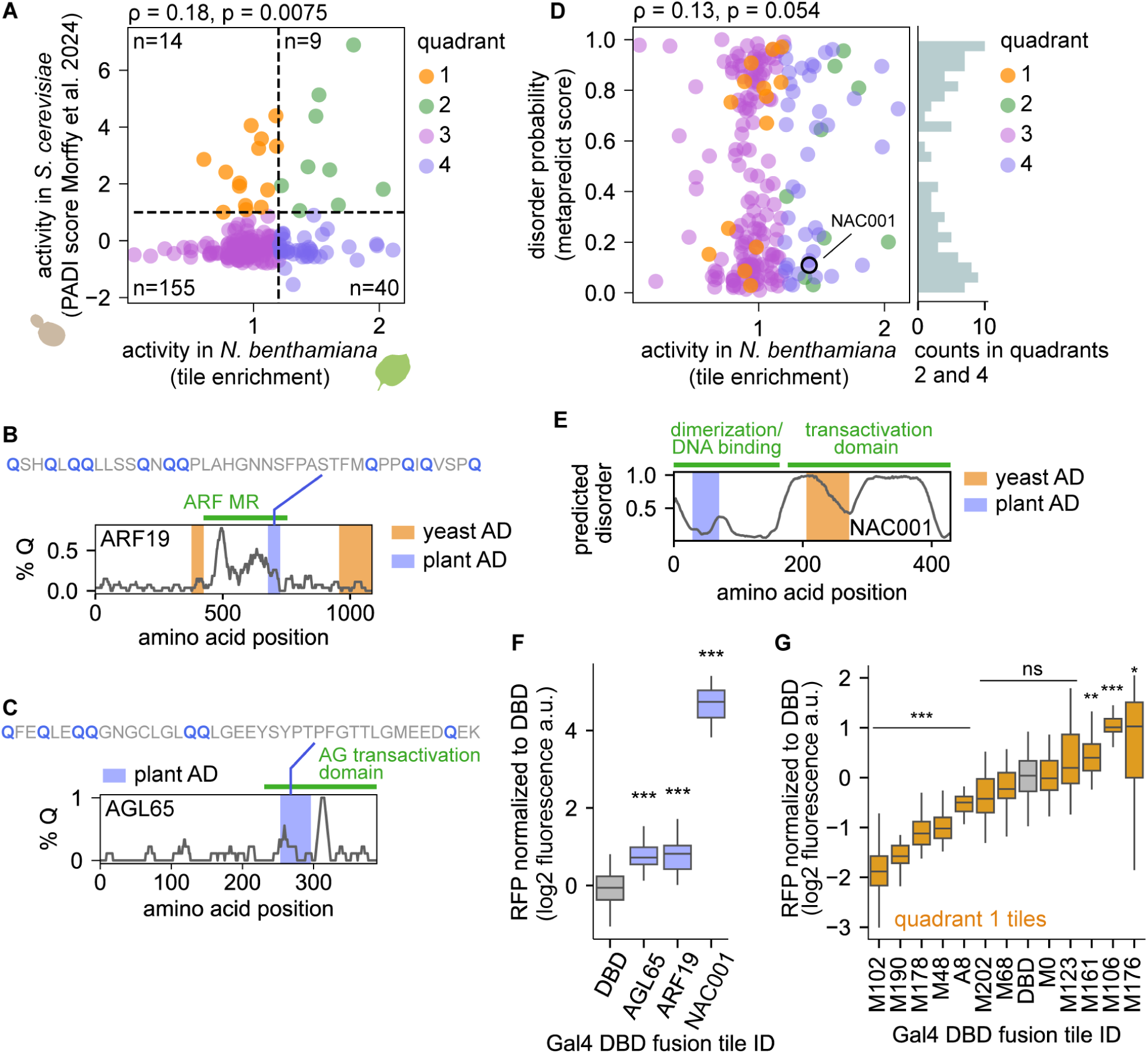
Systematic comparison of tiles from *Arabidopsis* TFs expressed in yeast and *N. benthamiana*. **(A)** Comparison of tile transcriptional activity between yeast and *N. benthamiana*. A set of peptide tiles from *Arabidopsis* TFs previously measured in yeast was tested in *N. benthamiana*. Scatterplot shows the activity of each tile in yeast (PADI score, y axis) and *N. benthamiana* (tile enrichment, x axis). The horizontal dashed line indicates the PADI threshold value used to classify tiles as activators in yeast. The vertical dashed line shows the enrichment value below which we never detected validated activator tiles in Fig. 3B. **(B)** Glutamine enrichment along the ARF19 TF protein shown as percentage of Q residues in a moving window of 20 aa. The green bar indicates the position of the ARF activation domain known as middle region (MR). Orange bars show the position of ADs according to the yeast assay. The blue bar shows the position of the AD in plants according to ENTRAP-seq. **(C)** Same as (B) for the ALG65 TF. No ADs were identified for this TF in yeast. **(D)** The degree of structural disorder of each tile is plotted against its ENTRAP-seq activity in *N. benthamiana*. The histogram on the right includes tiles likely to be activators in both yeast and *N. benthamiana* as well as *N. benthamiana* only. **(E)** Predicted disorder in the NAC001 TF. The green bar indicates the NAC dimerization domain. The blue bar shows the AD identified by ENTRAP-seq and the orange bar shows the position of the ADs active in yeast. **(F)** Validation of the tiles highlighted in blue in (B), (C), and (E) using a fluorescence reporter assay. **(G)** A random selection of tiles that were activators in yeast but de-enriched in tobacco (‘quadrant 1’ in (A)) tested in a fluorescence reporter assay against the Gal4DBD alone as a control. In (F) and (G) * p < 0.05, ** p < 0.01, *** p < 0.001, ns not significant in a two-sided Welch’s t-test against Gal4DBD with a Bonferroni multiple comparisons correction. In (F) and (G) box plots show the median ± IQR, whiskers correspond to minimum and maximum values excluding outliers.

An alternative hypothesis for why ADs can be active in plants but not in yeast is that these tiles are not ‘bona fide’ ADs, namely, peptides in intrinsically disordered regions that are sufficient to directly recruit the core transcriptional machinery to the promoter. Instead, these tiles may code for folded protein domains that bind plant-specific TFs present in N. benthamiana that in turn contain canonical ADs that activate transcription in yeast. To explore this possibility, we calculated the predicted disorder of tiles in the library and compared this prediction with their ENTRAP-seq enrichment. We found these two metrics to be completely uncorrelated (Spearman ⍴ = 0.13, p = 0.054) and tiles that are active in plants (enrichment >1.2) are equally likely to be structured as to be disordered (Fig. 5D). Thus, activator tiles with a plant-specific bias do not necessarily form folded domains. However, among highly structured tiles we found cases that do fit our initial hypothesis. For example, a tile in the NAC001 TF containing part of a structured domain involved in homo- and hetero-dimerization within NAC family TFs was found to be highly active in plants but not in yeast (Fig. 5E)^54^. The NAC TF family is plant-specific and contains acidic ADs in their C-terminal half that are activators in yeast^55^ (Fig. 5E). Thus, in *N. benthamiana*, this NAC001 tile might simply recruit endogenous NAC activators which, in turn, contain ADs that are active in both *N. benthamiana* and yeast. In yeast however, this NAC001 tile may be incapable of activating the reporter because this organism lacks interacting NAC partners.

We confirmed that the polyQ ADs of ARF19 and AGL65 as well as the NAC001 AD activated the reporter in our fluorescence plate reader assay (Fig. 5F). In this comparative ENTRAP-seq experiment, we found 14 tiles that had a PADI score > 1 and an enrichment value < 1.2, corresponding to likely yeast-only activators (labeled ‘quadrant 1’ in Fig. 5A). To test if these plant-derived sequences correspond to ADs that activate transcription in yeast but not in plants, we individually measured 12/14 of these tiles. Most (9/12) of these tiles did not activate transcription in *N. benthamiana* in a fluorescence reporter assay, in fact, 5/12 were repressors, consistent with their depletion from the ENTRAP-seq pulldown (Fig. 5G). These results are consistent with the notion that plant-specific as well as yeast-specific AD tiles are not uncommon. The stark differences in the rank-order of activity between the two systems for most tiles suggests that there are fundamental differences in transcriptional activation mechanisms between these two models, although other explanations are conceivable as well. For example, the reporter in the yeast assay is chromosomally integrated while the ENTRAP-seq reporter is transiently expressed. To better understand the degree of conservation of ADs between these two model eukaryotes, future studies will be necessary to leverage ENTRAP-seq to measure all tiles across all *Arabidopsis* TFs in an unbiased and systematic manner. Finally, these findings show that it is wise to exercise caution when interpreting tile MPRA data of plant proteins tested in yeast as both false positives and false negatives can arise when using one model system to represent the biology of another.

### Rationally tuning the activity of a plant TF

One of the implicit assumptions of AD recruitment assays is that TFs are largely modular^56^. If this is the case, it may be possible to study and engineer plant TFs in high throughput by isolating, mutating, and screening their ADs for altered functionality. Because ENTRAP-seq can reliably detect quantitative differences in activation strength we hypothesized that it could be used to screen for mutant AD variants displaying a continuous spectrum of activity. To test this idea, we leveraged ENTRAP-seq to characterize a mutant library of the *Arabidopsis* activator TF CONSTANS (CO) with the goal of creating an allelic series containing weaker and stronger activators than wild type as well as alleles that switch CO from activation to repression of target genes. Natural variation in CO and its homologs is associated with flowering time, a key agronomic trait^57,58^. Although *Arabidopsis* CO appears to act exclusively as a transcriptional activator, related CO-like genes in this and other species can act as repressors, suggesting that it is feasible to switch the CO regulatory role via mutations^59–61^. In agreement with the role of CO in its native context and previous reports, transient expression in *N. benthamiana* of a fusion between CO and a blue fluorescent protein (BFP-CO) strongly activates expression of the FT promoter (1.9 kb upstream of the ATG, hereafter FTpro) as measured by a GFP reporter assay (Fig. 6A)^62^.

**Figure 6:**
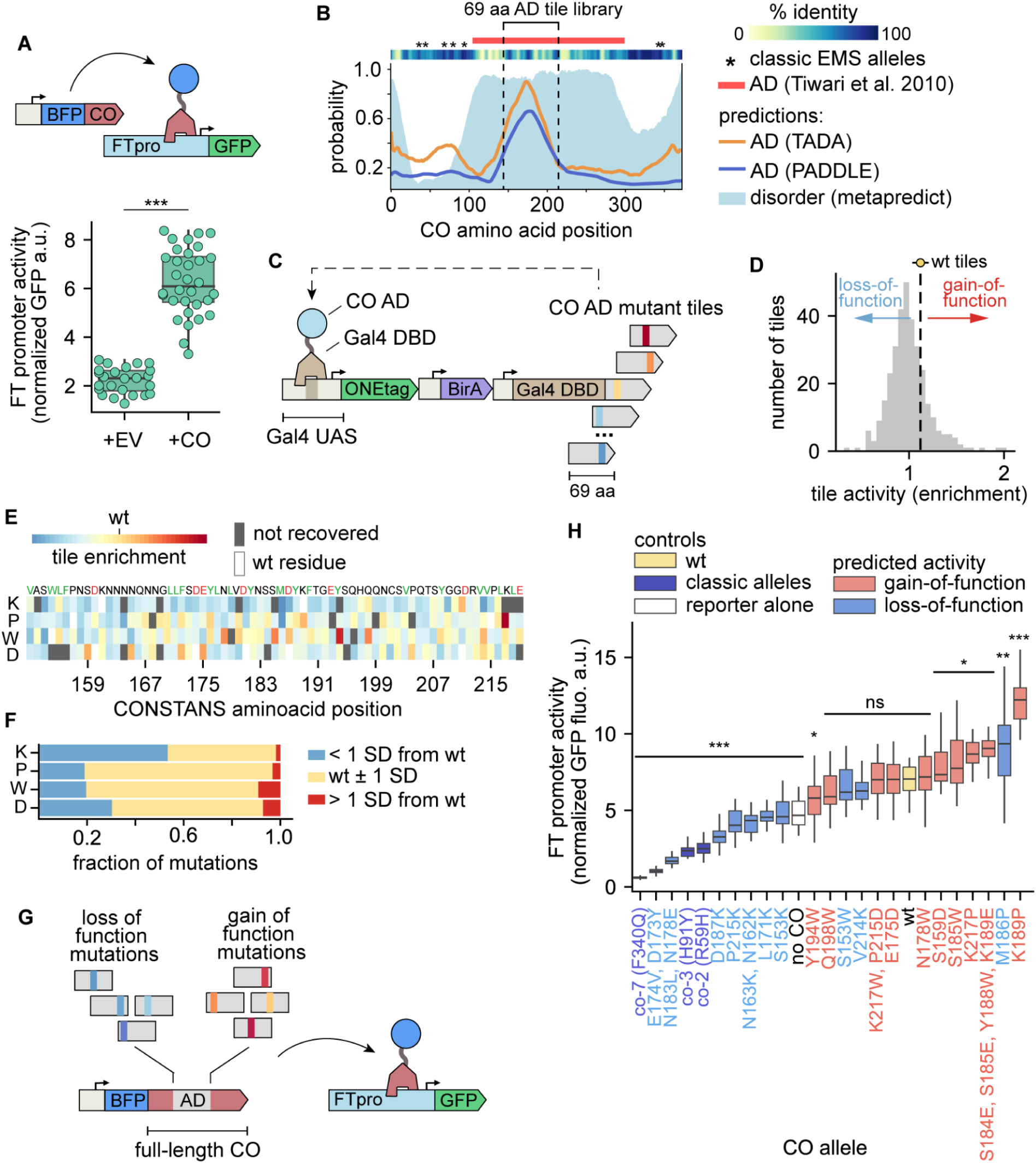
Large scale screening of mutant activation domains to create an allelic series of the CO transcription factor. **(A)** Transactivation assay for CO function in *N. benthamiana*. A strain carrying GFP driven by the *Arabidopsis* FT promoter (FTpro) was infiltrated with a strain carrying pUBQ-BFP-CO or an empty vector (EV). pUBQ-mCherry was used as a normalization. **(B)** Location of the CO activation domain (AD). Plot shows the probability of residues being part of an acidic AD according to the PADDLE and TADA as well as probability of being a disordered region according to metapredict. Bars on top show the empirically determined AD. The blue box indicates the 69 aa region used to build a tile library. **(C)** Screening of CO the AD tile library using nuclei enrichment. Mutant tiles are fused to the Gal4 DBD and used to drive the expression of the magnetic sorting reporter. **(D)** Distribution of tile enrichment values. The dashed line shows the mean enrichment of 6 wt tiles. **(E)** Tile activity as a function of biochemical properties. Each dot corresponds to one tile. The dashed red lines are linear fits. Shown on top is the correlation coefficient (R2) and the p-value of the linear regression. **(F)** Heatmap showing the activity of tiles carrying single aa substitutions. **(G)** Transactivation assay to test the effect of mutations in the CO AD in the context of the full-length protein. Mutations identified in (C)-(F) as being weaker (loss-of-function) or stronger (gain-of-function) than the wt AD were introduced into the CO coding sequence and used to drive expression of a FTpro GFP reporter. **(H)** FT promoter activity in the presence of various CO mutant alleles. Predicted activity is based on the enrichment of the corresponding AD tile with respect to the wt tile. In (A) and (H) box plots show the median ± IQR, bars correspond to minimum and maximum values excluding outliers. *** p < 0.001, ** p < 0.01, * p < 0.05, n.s not significantly in a Welch’s t-test comparison against WT with a Bonferroni multiple comparison correction.

CO contains a poorly conserved, highly disordered middle region that is sufficient for transactivation in a Gal4DBD recruitment assay (Fig. 6B)^62^. Although multiple CO alleles have been identified in random mutagenesis screens, all lie outside this middle region and are loss-of-function (Fig. 6B)^63^. To better define the position of the CO AD within this middle region, we leveraged PADDLE as well as TADA, another model trained on datasets from yeast-based screens that can predict acidic ADs^13,15^. Both models have a high degree of agreement and point to a region around aa position 175 as the putative CO AD (Fig. 6B). We therefore selected a 69 aa window in this region as the putative AD (Fig. 6B). To explore the sequence space of the CO AD, we generated a library of 350 tiles carrying 1-4 aa substitutions and including the wild-type AD as a control. The substitutions were rationally designed to either increase or decrease AD strength based on features important for acidic AD function in yeast and animal cells, namely 1) depletion of positive residues (R and K), 2) enrichment of negative residues (D and E), 3) enrichment of hydrophobic residues (particularly aromatics and leucine), 4) depletion of continuous stretches of hydrophobic residues, and 5) lack of stable secondary structure^6,13,64,65^. Following the same rationale, we also mutated every residue to K, P, W, and D. As internal controls we included 5 wt tiles. Finally, we introduced 10 random synonymous mutations in each tile in the library to better distinguish between highly similar tiles from sequencing. Following the same strategy used for our previous experiments, we cloned these tiles as Gal4DBD fusions in our magnetic sorting reporter and performed an ENTRAP-seq assay (Fig. 6C).

The enrichment of the CO AD tiles spanned a range of values comparable to other libraries and the enrichment of WT tiles was highly reproducible (1.12 ± 0.02). Perhaps not surprisingly, most tiles were less enriched than WT, corresponding to potential loss-of-function mutants (Fig. 6D). We found a moderate agreement between tile activity as measured by ENTRAP-seq and the predictions from PADDLE and TADA (Spearman ⍴ = 0.38 and 0.4, respectively) although the two models largely agreed with one another (Pearson r = 0.89) (Fig. S15). This suggests that models trained in plants could be exploited in the future to guide the design of mutant plant AD screens. Comparing the activity of each tile to its biochemical properties showed that, as expected for an acidic AD, negative charge and the number of F or W residues have a significant correlation with activity (Spearman ⍴ = -0.38 and 0.24, respectively, Fig. S16). Throughout the AD, mutation of residues to K tended to decrease the activity and most of the strong loss of function AD alleles are K substitutions (Fig. 6E, F). The majority of the gain-of-function single mutants correspond to W or D substitutions (Fig. 6E, F). Mutating residues to P, which severely disrupts protein secondary structure, rarely had strong effects (Fig. 6E, F). These results are consistent with the CO AD behaving like a classic intrinsically disordered acidic AD and highlights the power of combining predictive models with semi-rational mutagenesis guided by known AD compositional biases.

To test the effect of these mutations in the context of the full-length CO protein, we chose 10 putative loss-of-function and gain-of-function alleles among the tiles at either end of the enrichment value distribution. We then introduced the coding mutations from each of these tiles into the full-length CO expression construct and tested their effect individually using the FTpro fluorescence reporter previously established (Fig. 6G). As controls we also cloned three previously isolated semi-dominant loss-of-function alleles corresponding to point mutations outside the AD identified in random mutagenesis screens. Out of 10 predicted loss-of-function CO mutants, 7 were significantly weaker activators than WT and 2 of these were as weak as classic alleles. The M186P mutant behaved as a gain of function allele, in contrast to its activity as a Gal4 DBD fusion tile (Fig. 6H). The EMS mutant controls as well as some of our isolated alleles switched CO activity from an activator into a repressor, consistent with the EMS mutants being semi-dominant in *Arabidopsis*^63^. Out of the 10 predicted gain-of-function mutant alleles, 5 were stronger activators than WT (Fig. 6H). Among the rest, 4 retained activator function but were not statistically stronger than WT and one putative gain of function allele (Y194W) was a slightly weaker activator than WT. We conducted an independent replicate of this whole experiment and obtained similar results (Pearson r = 0.92, Fig. S17). Collectively, these data demonstrate that tile activity in the context of an ENTRAP-seq assay can be predictive of tile function within the original TF protein environment. This allowed us to rapidly engineer a new allelic series of a plant transcription factor, including alleles that switch its function from activator to repressor.

## Discussion

The long timescales required to generate transgenic plants has historically been a limiting factor in efforts to systematically map the sequence-to-function protein landscape of transcriptional regulators and other classes of protens^66^. Compared to stable transgenic approaches, protoplast transformation and transient expression in *N. benthamiana* can substantially accelerate the pace of sequence characterization of gene variants *in planta*. These systems have recently been used as expression platforms for multiplexed reporter assays to measure the activity of tens of thousands non-coding variants at a time ^24,67^. However, these technologies remain limited to library elements acting *in cis* to the reporter, such as enhancers and promoters. Gene libraries acting *in trans* (e.g., gRNA and protein-coding libraries) represent a unique challenge because transgenes that regulate the same reporter can readily interfere with one another if expressed in the same cell. To solve this problem, it was first necessary to achieve single transformations, which was only very recently demonstrated for agroinfiltration^26,28^. The second missing piece was linking transgene activity to a measurable output, something that ENTRAP-seq achieves by biasing the nucleus pulldown probability by the level of reporter expression.

While methods like DAP-seq ^8^ and TARGET ^9^ have shed light on TF DNA-binding preferences and *cis*-regulatory landscapes, they do not capture how TF regulatory activity itself is encoded at the protein level. This gap is particularly critical given that many transcriptional regulatory domains — such as ADs — are intrinsically disordered and poorly conserved, making them difficult to identify through structure- and homology-based algorithms. Beyond mapping transcriptional circuitry, ENTRAP-seq provides a powerful framework for dissecting and engineering protein function. Its parallelized, library-based design is well-suited to systematic exploration of sequence-to-function relationships, akin to deep mutational scanning approaches that have revealed critical residues and domains in other systems^66^. By screening libraries of natural variants or synthetic constructs, ENTRAP-seq enables the direct measurement of how coding sequence variation impacts regulatory activity—an essential step in linking genotype to phenotype. This capability is particularly relevant for proteins involved in pathways that are not well conserved across eukaryotes and has broad implications for plant biology and crop improvement.

As a method to quantify reporter activity, ENTRAP-seq shows a remarkable level of agreement with traditional low throughput approaches. Indeed, combining all the cross-comparisons between enrichment and fluorescence reporter assays presented in this study reveals a tight correlation (Fig. S18, Spearman ⍴=0.83, N=497 comparisons in total). This meta-analysis also shows that our method to define activators based on enrichment is highly conservative: there are zero tiles that pass the enrichment cutoff but fail to activate the reporter in a fluorescence assay (Fig. S18). This lack of false positives at the expense of false negatives should be kept in mind when interpreting our AD identification results. The degree of enrichment is a quantitative metric of transcriptional strength, but this information is necessarily lost when collapsing this continuous quantitative metric into a binary. Although the absolute levels of enrichment of ENTRAP-seq (approximately 0.5-2.5X) are small compared to other enrichment-based MPRAs, the standard error of this method is also much smaller. Scaling the enrichment of ENTRAP-seq and a recently published enrichment-based cell culture MPRA ^22^ by their corresponding standard error shows that both methods have comparable dynamic ranges (Fig. S19). Together, these findings highlight the robustness, precision, and conservative reliability of ENTRAP-seq as a quantitative approach for identifying and characterizing transcriptional activators.

Recently, genome editing technologies have opened the door to generating TF alleles with quantitative differences in activity as a means to modulate quantitative crop phenotypes ^68^. These efforts have focused on editing *cis*-regulatory regions rather than TF coding sequences. We envision that coding mutations in TF regulatory domains may provide an alternative approach to engineer such allelic series. Compared to *cis*-regulatory mutations, coding mutations are more likely to conserve expression levels and spatiotemporal expression patterns. These mutations could be identified by combining ENTRAP-seq screening with predictive sequence-to-function models of TF activity. Because quantitative and qualitative aspects of plant transcription are not necessarily conserved in yeast, future models of plant TF activity will need to be trained on diverse plant-derived sequences expressed in plant cells. First generation yeast models such as PADDLE were based on 5-10K protein tiles, a scale that is within reach by combining a few ENTRAP-seq runs. Similarly, by combining several ENTRAP-seq runs, genome scale coverage of all TFs in a plant genome may also be feasible. Overall, ENTRAP-seq addresses a key technical gap in plant functional genomics by systematically linking protein sequence to transcriptional output in a native plant context, allowing for the functional annotation of regulatory domains at scale.

## Acknowledgments

First and foremost we would like to thank all members of the Shih lab. From weekly growing healthy plants to making *Agrobacterium* competent cells to discussing results, their support enabled this research. In particular, we are thankful to Niklas Hummel for introducing us to agroinfiltration and the study of TFs in plants and for sharing countless plasmids. Amy Lanclot and Maximo Kesselhaut provided insightful comments during the preparation of this manuscript. Anthony Sarkiss helped with infiltrations. We also thank a number of colleagues with whom we discussed preliminary data and ideas during the development of this project. We had productive and inspiring discussions with Max Staller, Roger Deal, Jose Miguel Alvarez, Patricia Lang, Hernan Garcia, Christine Queitsch, and George Coupland. A number of scientists provided critical technical feedback as well. Yu Tang and Julia Bailey-Serres shared constructs for nuclear envelope targeting, Adam Deutschbauer gave us advice for library cloning and sequencing and Rita Kuo and Noam Prywes guided us through the early stages of next-generation sequencing. Finally, we thank Shana McDevitt from QB3 Genomics, UC Berkeley, Berkeley, CA, RRID:SCR_022170 and Brian McCarthy from the UC Berkeley DNA sequencing facility for their advice and assistance. This work was part of the DOE Joint BioEnergy Institute (https://www.jbei.org) supported by the U. S. Department of Energy, Office of Science, Office of Biological and Environmental Research, through contract DE-AC02-05CH11231 between Lawrence Berkeley National Laboratory and the U.S. Department of Energy.

## Data, code, and sequencing availability

All code and models related to this study are publicly available on GitHub (github.com/shih-lab/magnets). Raw sequencing reads are publicly available through the NCBI SRA Database under BioProject accession PRJNA1254296.

## Author Contributions

P.M.S provided strategic scientific guidance and funding for this project. P.M.S, L.W, and S.A conceived the research and wrote the manuscript. L.W and S.A analysed the data. L.W performed statistical inference, modeling, and phylogenetic analyses. S.A, A.D, L.O, R.R, and S.O contributed biological materials. S.A performed experiments.

## Materials availability

All plasmids are available from the JBEI registry. The sequences of all plasmids used in this study are listed in table S4. The sequences of plasmids that were created for this study can be found in the public google drive folder: https://drive.google.com/drive/folders/1ogtAr0cA7BEVsjFaDrkzEIUccGWJD2VS?usp=drive_link

## Materials and Methods

### Molecular cloning

Except for dCas9 fusions, all validation and candidate tiles tested individually in fluorescence reporter assays were cloned in the pCambia1300-based backbone pSA676v2 containing 3 transgenes: 6xUAS-min35S::RFP-T2A-SlWIP2-C ONEtag, pUBQ10::BirA, and pMas::Gal4DBD-dropout. The dropout consists of a E.coli GFP reporter flanked by BsaI sites for negative GFP colony selection. Individual tiles were cloned in BsaI-digested pSA676v2 using Gibson assembly. Tiles fused to dCas9 were cloned into backbone pSA742 containing p35S::dCas9-dropout using Gibson assembly. The sequence of pSA676v2 and pSA742 as well as other plasmids used in this study can be found in the drive folder listed above. For library cloning see Materials and Methods: Library cloning.

### Live imaging of leaf tissues

Leaf samples 3 days post infiltration were mounted in water. All images were acquired in a Zeiss LSM 710 confocal microscope using a 63X oil objective. Channels were acquired sequentially using 20 z slices every 0.5 μm and 1024 x 1024 pixel image size. The zoom was 2.0 and the wavelengths were 568 nm excitation, 585-630 nm emission for RFP, and 488 nm excitation, 494-581 nm emission for GFP. The pinhole in both channels was set to 1 AU.

### Imaging and fluorescence quantification of isolated nuclei

Purified nuclei were mounted in NPB buffer in a hemocytometer and imaged using a Leica DM6B epifluorescence microscope. 8 images were acquired from each purification fraction. For each image, three z-stacks were acquired sequentially, one for each channel. Each z-stack consisted of 5 z-sections every 20 µm. The BFP nuclear marker was imaged with DAPI filter (excitation 350±50, emission 460±50), the ONEtag was imaged using the L5 filter (excitation 480±40, emission 527±30), and mCherry RFP was imaged using the TXR filter (excitation 560±20, emission 630 ±38). Images were acquired using a 5x dry objective for an area of 2.641 mm^2^ and 2048 x 2048 pixels (pixel size = 0.78 µm^2^). To avoid detector saturation, the laser power and the camera exposure were set to 17% and 600 ms, respectively, for all channels. Stacks were max-projected in z for all subsequent analyses. Representative BFP images were manually curated to label nuclei and background and these labels were used to train a random forest classifier. This model was then used to generate a binary segmentation of all BFP images. Only objects with an area between 40-400 µm^2^ were considered as true nuclei. Binary nuclear masks were superimposed on the GFP and RFP images to calculate the nuclear fluorescence in these channels. Nucleus fluorescence was calculated as the average background-subtracted pixel intensity of the 20% brightest pixels. The average pixel intensity outside a dilated nuclear mask was used as background.

### Library cloning

Single stranded oligo pools containing tiles of each library were obtained from Twist Biosciences. The oligos were flanked by a shared overhang sequence containing BsaI cut sites for cloning into the ENTRAP-seq backbone pSA676v2. The overhangs were 5’ ACGTCAGTGTG<u>GTCTCT</u>**AGGT** and TAA**GCTT**<u>AGAGAC</u>CAAGCGCTTA 3’ (BsaI sites underlined, cloning overhangs in bold). Oligo pool DNA was resuspended in 150 μl of water and used for a 50 μl PCR reaction using primers binding to the overhangs (tileLib-F and tileLib-R, table S5). The PCR reaction contained 1 μl of resuspended oligos as template, 2.5 μl of each primer at 10 uM, 19 μl of water, and 25 μl of 2x NEB Q5 high fidelity DNA polymerase master mix. The reactions were run using the standard conditions provided by the Q5 manufacturer using a melting temperature of 65C, an elongation time of 15 sec and 15 cycles. A control 10 μl reaction was run for 35 cycles. Both reactions were run side by side on a 1% agarose gel to confirm correct amplification. The PCR was purified using Zymoclean DNA clean and concentrator columns using two washes with 600 μl of wash buffer and eluting in 30 μl of water. The dsDNA amplified oligo pool was then digested overnight at 37C using the following conditions: 13 μl of water, 30 μl of purified PCR, 5 μl of NEB 10X HF buffer, 2 μl of NEB BsaI-HF. The digestion was purified using Zymoclean DNA clean and concentrator columns using two washes with 600 μl of wash buffer and eluting in 20 μl of water. For the backbone, approximately 10 μg of pSA676v2 DNA were obtained from a standard miniprep. The plasmid was digested overnight at 37C with NEB BsaI-HF using the following reaction conditions: 50 μl plasmid miniprep DNA, 3 μl of restriction enzyme, 8 μl of 10X HF buffer, 19 μl of water. The entire digestion was run on a 1% agarose gel and the band corresponding to the backbone was purified using the Zymoclean gel DNA recovery kit using two washes with 600 μl of wash buffer and eluting in 20 μl. The digested dsDNA library was ligated into digested pSA676v2 using NEB T7 ligase. 500 ng of backbone (approximately 6 μl) and enough insert to achieve a ratio of 3:1 insert:backbone were added to 120 μl final reaction volume ensuring that the total DNA concentration did not exceed 10ng/μl. The reaction was run using 3 cycles of 2 hr at 22C and 4 hr at 16C. The ligations were column-purified and eluted in 15 μl of water.

### Library transformation into bacteria

One tube of *E. coli* SIG10 MAX electrocompetent cells (Millipore Sigma CMC0004) was electroporated with 2 μl of ligated library and recovered in LB at 37C for 1 hr. Following recovery, 1 μl of cells was plated in 50 μg/ml Kanamycin LB plates in triplicate to assess the number of colony forming units (CFUs) per transformation. The minimum number of CFUs used was 10 times the library size. The rest of the culture was inoculated in 50 ml of 50 μg/ml Kanamycin LB and grown overnight at 37C. A miniprep was run on 5 ml of the overnight culture to obtain plasmid DNA for *Agrobacterium* transformation. For each library, *Agrobacterium* GV3101::pMP90 was electroporated using 10 μl of competent cells, 2 μl of library miniprep (approximately 500 ng of DNA) and 88 μl of water in each cuvette. Cells were recovered in 1 ml of LB at 28°C for 2 hr. 5 such transformations per library were used to increase the number of colony forming units (CFUs). Following recovery, 1 μl of cells was plated in LB plates containing 50 μg/ml Rifampicin, 50 μg/ml Kanamycin, and 30 μg/ml Gentamicin in triplicate to assess the number of CFUs per transformation. The minimum number of *Agrobacterium* CFUs used was 10 times the library size. Recovered *Agrobacterium* cells were inoculated in 100 ml of LB containing the same antibiotic concentrations and grown for 2 days at 30°C shaking. Glycerol cryostocks were prepared from these cultures.

### Plant growth conditions

*Nicotiana benthamiana* was grown in a controlled environment room at 25°C of temperature, 60% humidity, 16/8 hr light/dark cycle, and a light intensity of 120 μmol/m^2^s. Prior to agroinfiltration, plants were grown for 29 days (sowing at day 0) in individual pots using Sunshine Mix #4 soil (Sungro) supplemented with 499 Osmocote 14-14-14 fertilizer (ICL) at 5mL/L. Immediately after agroinfiltration plants were returned to the growth room for 3 days until sampling.

### *Agrobacterium* growth conditions

For fluorescence plate reader assays *Agrobacterium* glycerol stocks were streaked onto LB agar plates supplemented with antibiotics: 50 μg/ml Rifampicin, 50 μg/ml Kanamycin, and 30 μg/ml Gentamicin. The day prior to infiltration, individual colonies were inoculated into liquid LB medium containing the same antibiotics and incubated overnight at 30°C, shaking. On the day of infiltration, cultures were diluted 1:10 into LB with identical antibiotic concentrations and incubated further under the same conditions for 3 hours or until an OD (optical density at 600 nm) of ∼1.0 was reached. For gene libraries, transformed cells previously recovered for 2 hr in LB without antibiotics were transferred to 100 ml of LB containing the same antibiotic concentrations listed above. Two days later, the OD of this culture was measured and the culture was diluted 1:10 if the OD was higher than 1.0 prior to infiltration. Alternatively, approximately 100 μl of library glycerol cryostocks were inoculated into LB containing antibiotics the day before infiltration. The day of infiltration, the culture was diluted 1:10 and grown until reaching an OD of ∼1.0.

### Agroinfiltration

*Agrobacterium* liquid cultures with an OD between 0.5 and 1.0 were pelleted by centrifugation at 4,000 G for approximately 10 minutes and resuspended in an equal volume of infiltration buffer (10 mM MES, pH 5.6; 10 mM MgCl₂; 150 μM Acetosyringone). The resuspended cultures were incubated with shaking at room temperature for 1 hour before OD measurements were taken. Next, a 1:5 dilution of each culture was prepared in infiltration buffer, and the OD of this dilution was determined using a spectrophotometer. Final OD dilutions for infiltration were then made using the same buffer. The mixtures were infiltrated into the 6th and 7th leaves (counting upward, with cotyledons considered as leaves 1 and 2) within 1 hour of preparation. For fluorescence reporter assays 3 mixes were infiltrated per leaf at an OD of 0.2 each (total bacterial OD of 0.6). For nucleus purification infiltrations the culture was diluted to OD 0.0025.

### Fluorescence plate reader reporter assays

For leaf fluorescence measurements of transcriptional reporters, 4 6mm leaf disks were cut from each agroinfiltrated leaf using a single-hole puncher and deposited on top of 350µL of tap water in a clear-bottom black 96-well plate. All reporter constructs were cloned in the ENTRAP-seq construct backbone (pSA676v2). The DBD-only control contains a stop codon instead of a tile (pSA676v2s). For normalization a pMAS::BFP strain was coinfiltrated (pSA639). Both strains were diluted to OD 0.4 and mixed in equal ratios to a final OD of 0.2 each. Three days post infiltration the plates were read on a BioTek Synergy H1 plate reader using 488 nm excitation and 520 nm emission for GFP, 587 nm excitation and 615 nm emission for RFP, 400 nm excitation and 454 nm emission for BFP. The normalized reporter activity reported throughout this work corresponds to a double normalization to BFP and to Gal4DBD alone. First, the RFP fluorescence of each leaf punch was divided by its BFP fluorescence. Next, this value was divided by the mean RFP/BFP fluorescence of Gal4DBD.

### Nucleus Purification Buffer (NPB and NPBt)

NPB was prepared following the recipe from Wang & Deal 2015: 20 mM MOPS pH 7, 40 mM NaCl, 90 mM KCl, 2 mM EDTA pH 8, 0.5 mM EGTA pH 8, 0.5 mM spermidine, 0.2 mM spermine, 1 complete protease inhibitor tablet per 100 ml. A 5x stock solution was prepared adding everything except for fresh ingredients: spermidine, spermine and protease inhibitors. The fresh ingredients were added to the 1x solution up to one day prior to nucleus pulldown. NPB with triton (NPBt) was prepared following the same recipe and protocol except that 0.1 % (v/v) Triton X-100 was added.

### Preparing tissue for nucleus sorting

*Agrobacterium* cultures containing gene libraries were diluted to a final OD of 0.0025 in IB and infiltrated into 8-10 4-week old *N. benthamiana* plants (see Materials and Methods: Agroinfiltration). The full area of true leaves 8-10 of each plant was infiltrated. Three days post infiltration, leaves were harvested and the mid vein was removed. A ∼5mm perimeter around the edge of the leaves was discarded as well. Ten grams of leaf material was then chopped into ∼1cm^2^ pieces using a razor blade. The chopped leaves were submerged in 50 ml of nucleus purification buffer (NPB) in a beaker and placed in a vacuum chamber for 10 min to infiltrate NPB into the tissue (see Materials and Methods: Nucleus Purification Buffer). Further rounds of vacuum filtration were applied until the tissue was completely soaked in NPB. Next, the tissue was washed 3 times with tap water and tapped dry with paper towels. Finally, the leaf material was flash frozen in liquid nitrogen, ground into a very fine powder using mortar and pestle and stored at -80° C.

### Nucleus pulldown

Frozen ground tissue (see Materials and Methods: Preparing tissue for nucleus sorting) was resuspended in 100 ml of ice-cold NPB and passed through a 70 μm strainer (Corning CLS431751) into 2 50 ml tubes to separate nuclei from cells and other larger leaf fragments, keeping the material on ice at all times. This filtrate was centrifuged at 500 G for 10 minutes at 4°C. The pellets from both tubes were combined by resuspending them in 20 ml of NPB. The resuspension was passed through a 100 μm strainer (Fischer Scientific 07000221) to remove remaining pieces of tissue. In parallel, 100 μl of Dynabeads MyOne streptavidin beads C1 (Thermo Fischer 65001) were washed by resuspending them in 1.5 ml of NPB, separating them in a magnetic rack and resuspending in 100 μl of NPB. The washed dynabeads were then added to the resuspended nuclei and the mix was incubated for 30 minutes, shaking gently in a cold room. The following steps were carried out in a cold room. The nuclei were diluted in NPBt to 100 ml final split into 2 50 ml tubes. Each tube was then attached to 4 neodymium magnets (LOVIMAG # B0CBTSFFSB) using an elastic band (See Fig. S20 for details) and placed in this setup for 30 minutes to pull down nuclei. The flow-through of this pulldown (i.e., the supernatant) was carefully removed and centrifuged for 10 min at 500 G, resuspended in 25 ml of NPB per tube and centrifuged again at 500 G for 10 min to obtain a pellet corresponding to the flow-through sample. The pulldown from the first magnetic separation was washed twice by resuspending it in 50 ml of NPBt and placing it in the magnetic rack for 20 min. The washed nuclei were next resuspended in 15 ml of NPBt in a 15 ml tube and placed in the magnetic rack again for 20 min. The pellet from this separation was resuspended in 3 ml of NPB and 10 μl of this sample was taken for imaging. The rest of the sample was centrifuged at 1000 G for 10 min to obtain a pellet corresponding to the pulldown sample. The pellets corresponding to the flowthrough and pulldown samples were resuspended in 1 mL and 200 μl of CTAB, respectively (100 mM Tris-Cl pH 8, 20 mM EDTA pH 8, 1.4 M NaCl, 2% (w/v) cetyltrimethyl ammonium bromide, 1% PVP 40,000). These samples were stored at -20°C until DNA extraction.

### Nucleus DNA extraction

Purification of DNA from the flowthrough and pulldown samples was carried out based on a previously published protocol^69^. CTAB-resuspended pellets were incubated at 65°C for 30 minutes vortexing every 10 minutes. A volume approximately equal to the pellet volume of phenol:chloroform:isoamyl alcohol 25:24:1 v/v was added per tube and tubes were vortexed thoroughly. Samples were centrifuged at 20K G for 2 min at room temperature. The top aqueous layer was removed and mixed with an equal volume of CTAB in a new tube. Samples were vortexed and incubated at 65°C for 20 minutes. A volume of chloroform:isoamyl alcohol 96:4 v/v equal to the volume of CTAB added in the last step was added to each tube. Samples were vigorously vortexed and centrifuged at 20K G at room temperature for 2 minutes. The top aqueous layer was transferred to a new tube and 70% of this volume of isopropanol was gently added to the wall of each tube. Samples were mixed gently by inverting 5-10 times and placed and -20°C for 20 min. The DNA was pelleted by centrifugation at 20K G for 15 min. The supernatant was removed and the pellets were washed with ice-cold 70% ethanol. Ethanol was removed and the pellets were air dried on a bench top prior to resuspending them in either 54 μl (pulldown) or 108 μl (flow-through) of water.

### Sequencing library preparation

For each sample, a 480 μl PCR reaction was prepared containing: 240 μl of Q5 2x master mix, 168 μl of water, 24 μl of nucleus DNA extraction, and 24 μl of a 10 μM stock of each primer. Primers were designed to anneal to regions immediately adjacent to the tile sequences and contained Nextera-style sequencing adapters and 8 bp indexes. Throughout this work we used the same number of PCR cycles for all samples, which was determined as follows. We ran qPCR reactions using the pulldown and flow-through DNA samples of 2 replicates of the viral 250 tile library, using the same set of primers used for preparing sequencing libraries. The qPCR Ct value of these reactions was 22-23 cycles. We added 2 more cycles to obtain extra DNA for sequencing, resulting in 25 cycles used for library preparation PCR. The library PCR were run on a 2% agarose gel and bands of the expected size were excised and purified using a Zymo gel purification column following manufacturer instructions. Libraries were eluted in 50 μl of water and run in a fragment analyzer to confirm that the size distribution matched the expected amplicon size. Whenever necessary libraries were size selected using SPRI bead cleanup. Samples from the 26 control strains experiments shown in Figure 2 were PCR amplified using the same protocol except that primers did not contain any extensions. Adapters were added to the purified PCR products using the Nanopore Native Barcoding Kit 96 V14 (SQK-NBD114.96).

### DNA sequencing

The libraries prepared from the control set of 26 mixed strains shown in Figure 2 were sequenced using the entire whole flow cell of a Nanopore minION for each replicate. The rest of the samples were run on an Illumina NextSeq 2000 for 100 cycles using 30% PhiX and single-end reads covering the 5’ end of the sequencing library amplicon.

### Mapping reads to tiles

The 2FAST2Q^70^ package version 2.6.0 was run using the following parameters. Feature length = 76 bp, allowed mismatches = 2, minimal Phred score = 20, upstream search sequence = TGGATCTGGAGGT, mismatches in upstream sequence = 0, minimal upstream sequence Phred score = 30.

### Calculation of tile enrichment

For each tile, the number of mapped reads in the pulldown sample was divided by the total number of reads mapped to all the library tiles in this sample to obtain a number corresponding to the proportion of pulldown reads mapping to each specific tile. The same calculation was performed for each tile in the flowthrough sample. The ratio between these two values corresponds to the tile enrichment.

### Calculation of tile coverage

We define tile coverage as the percentage of all reads in both pulldown and flowthrough samples that map to a given tile. To obtain this percentage the number of mapped reads in the pulldown and flowthrough samples are added and the reads that match each tile are divided by the total number of reads.

### Designing viral libraries

A total of 1,495 plant virus genomes were downloaded from NCBI, yielding 5,607 unique proteins. These proteins were clustered at 90% sequence identity using CD-HIT ^71^, resulting in 4,537 representative proteins. Each protein was fragmented into 53 aa tiles with a 10 aa stride, generating 190,520 tiles. Tile size was kept to 53 aa to match the tile size in the dataset used to train the PADDLE AD prediction model ^14^. This size is large enough to capture common ADs while minimizing PCR bias due to long amplicon size ^3,4^. PADDLE was used to perform inference on each tile. To construct a diverse subset for experimental screening, the top 2,000 tiles with the highest predicted z-score were binary encoded, and a greedy selection algorithm was applied to maximize the pairwise Hamming distance between selected encodings. This approach prioritized sequences with the most dissimilar binary representations, thereby enhancing diversity in the DNA library.

### Machine-based predictions of activation domains

In addition to PADDLE, TADA was used to predict transactivation strength, and Metapredict^72^ was used to identify disorder regions in proteins. Because PADDLE and TADA use different input sizes (53 aa and 40 aa, respectively), CONSTANS and CO variants were tiled with a 1 aa stride, and tile-level predictions were aggregated into residue-level profiles by averaging the scores of all overlapping tiles at each position. Residues not covered by any tile were assigned a score of zero. PADDLE and TADA predictions were independently min–max normalized to scale scores between 0 and 1 for comparison between models.

### Calibration of ENTRAP-seq from fluorescence data

To model the relationship between ENTRAP-seq and fluorescence measurements, we fit a sigmoid function to the enrichment ratios as a function of logRFP for control tiles. The standard deviation of residuals from the fitted curve was used to estimate confidence bounds. We defined the activation threshold as the upper confidence bound at the midpoint of the sigmoidal fit.

## Supplementary Figures

**Figure S1:**
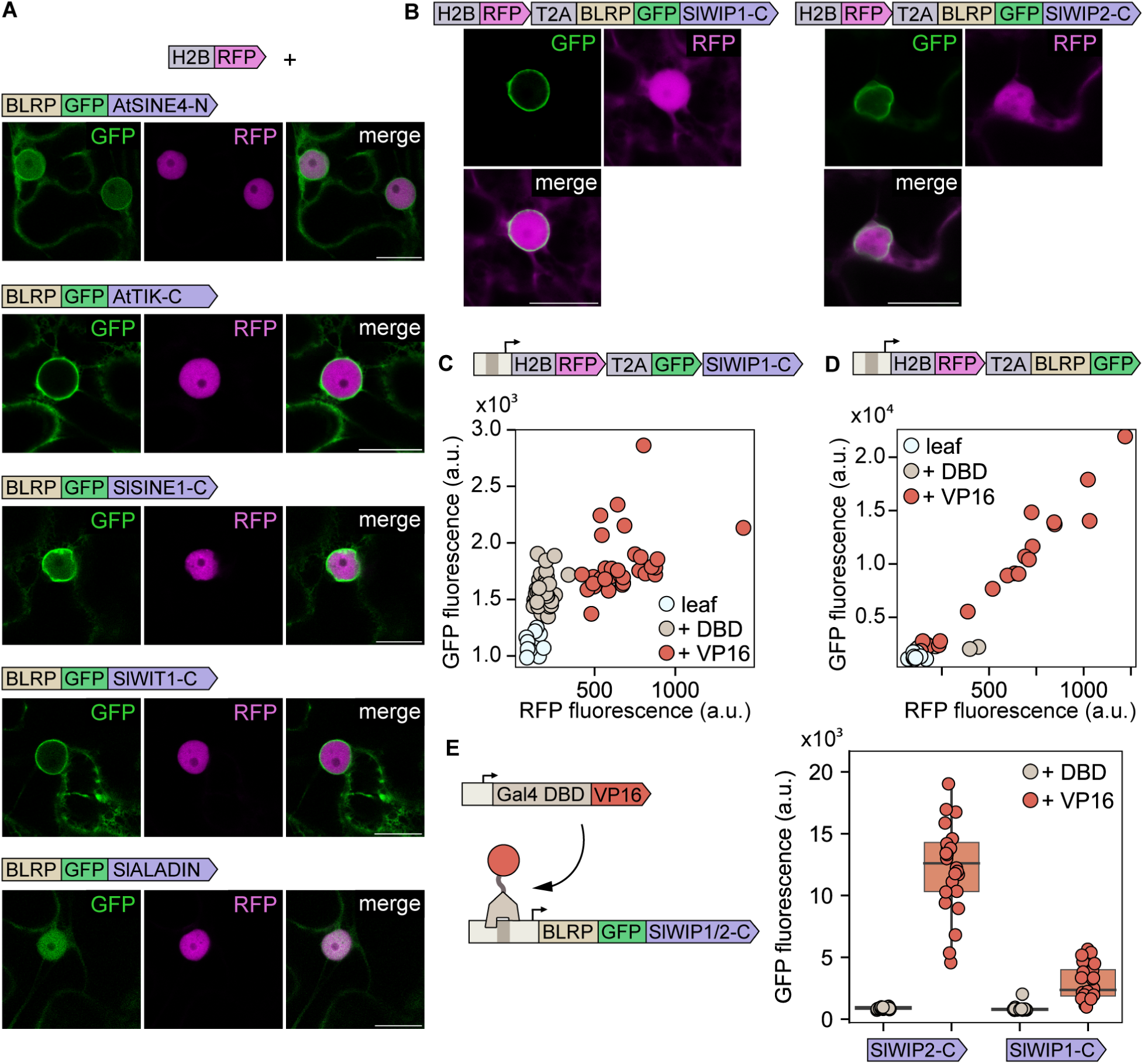
Additional ONEtag optimization details. **(A)** Fluorescence microscopy images showing the subcellular localization of ONEtag variants tagged with GFP and coexpressed with an RFP nuclear marker. From top to bottom the corresponding plasmid names are pSA266, pSA267, pSA268, pSA270, pSA272. **(B)** Cells expressing the SlWIP1-C and SlWIP2-C ONEtag fused to RFP by a self-cleaving T2A peptide (plasmids pSA271 and pSA91, respectively). **(C)** Top: schematic of the construct. BLRP was removed from the construct shown in Fig. 1C and the responsiveness of this reporter to VP16 was tested in a fluorescence plate reader. **(D)** Top: schematic of the construct. SlWIP1-C was removed from the construct shown in Fig. 1C and the responsiveness of this reporter to VP16 was tested in a fluorescence plate reader. In (C) and (D) each circle corresponds to one leaf punch. **(E)** Left: schematic of the experiment. The RFP-T2A tag was removed from both the SlWIP1 and SlWIP2 constructs shown in Fig. 1C and their responsiveness to VP16 was tested in a fluorescence plate reader (plasmids SA752 and pSA753, respectively). Right: leaf GFP fluorescence with or without VP16 driving reporter expression. Box plots show the median ± IQR, whiskers correspond to minimum and maximum values.

**Figure S2:**
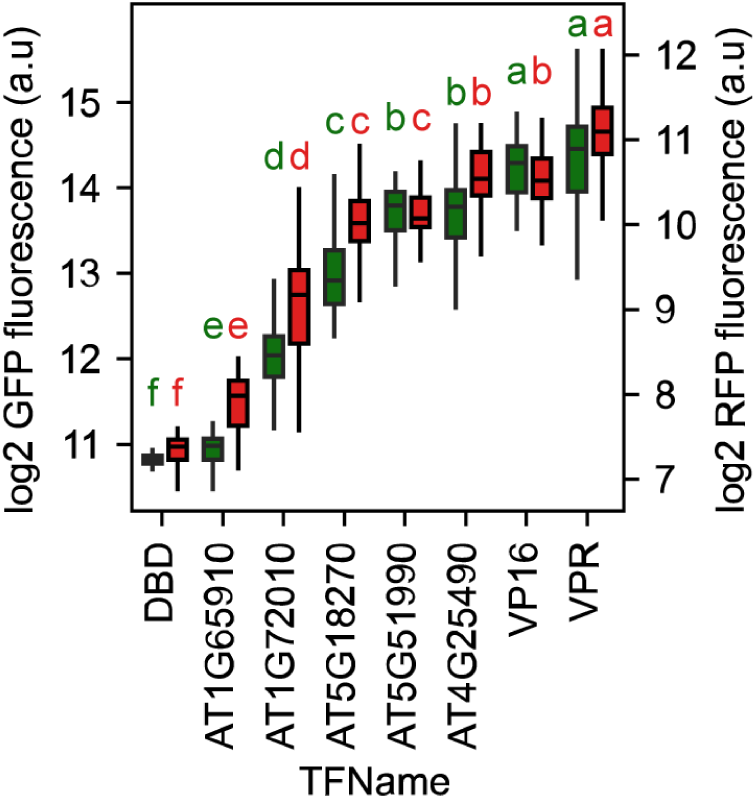
Related to Figure 1G. Sensitivity of RFP-T2A-SlWIP2-C as a fluorescence reporter. Fluorescence of leaves infiltrated with the 6xUAS-min35S-RFP-T2A-SlWIP2-C reporter in combination with Gal4DBD fusions with different TFs or activation domains (same data as in Fig. 1G). Box plots show median ± IQR, whiskers correspond to minimum and maximum values. The letters on top of each box represent statistically significant groups based on a post-hoc Tuckey Honestly Significant Difference (HSD) test performed separately for GFP and RFP.

**Figure S3:**
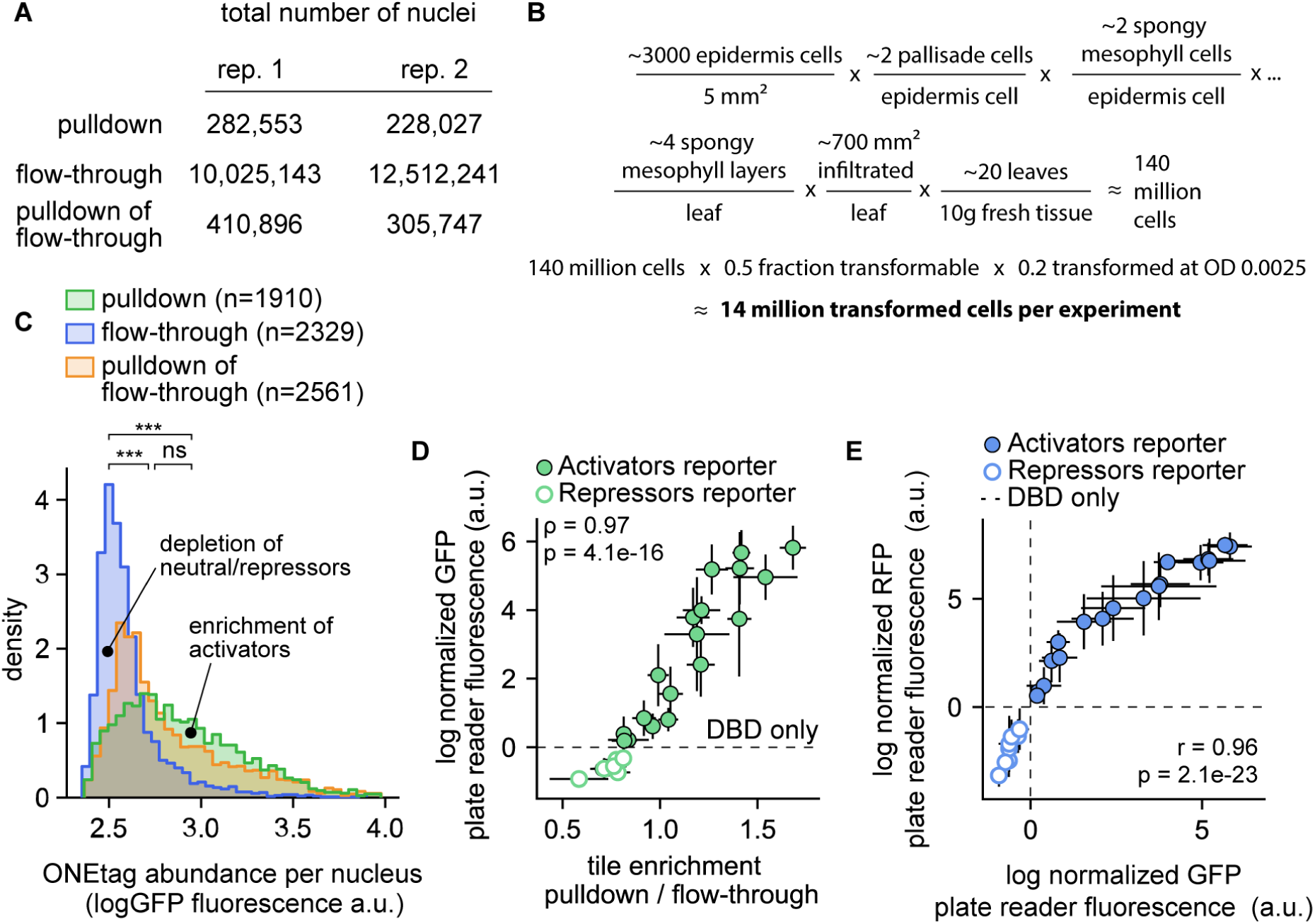
Validation details of ENTRAP-seq for quantitative measurements of reporter expression. **(A)** Total number of nuclei obtained in each purification fraction in the experiment from Fig. 2C (rep.1) and Fig. S2A (rep.2). **(B)** Order of magnitude estimate of the number of cells transformed by a strain carrying the ONEtag based on the numbers obtained from Alamos et al. 2025^26^. **(C)** Biological replicate of the experiment shown in Fig. 2C. Shown are the distributions of ONEtag expression levels in nuclei from different purification fractions. The histograms are normalized to have the same area. Nuclei with low GFP levels are enriched in the flow-through and there is a progressive enrichment in the pulldown with increasing GFP expression. *** p<0.001, ns distributions not significantly different using Mann–Whitney U test. **(D)** Extended data for the experiment from Fig. 2E,F. X-axis: transcriptional activity of each tile based on ENTRAP-seq (mean ±SD enrichment of each tile T-DNA in the nuclei pulldown fraction compared to the flow-through sample). Y-axis: mean ± 1 s.d. GFP expression measured for each tile individually using a fluorescence plate reader. **(E)** Extended data for the experiment from Fig. 2E,F. Mean ± 1 s.d. GFP (x-axis) and RFP (y-axis) fluorescence for each control tile. N for nuclei pulldown and sequencing = 4 biological replicates. N for plate reader assays = 24 (3 plants with 8 leaf punches per plant). ⍴ = Spearman correlation coefficient, r = Pearson’s correlation coefficient.

**Fig. S4.**
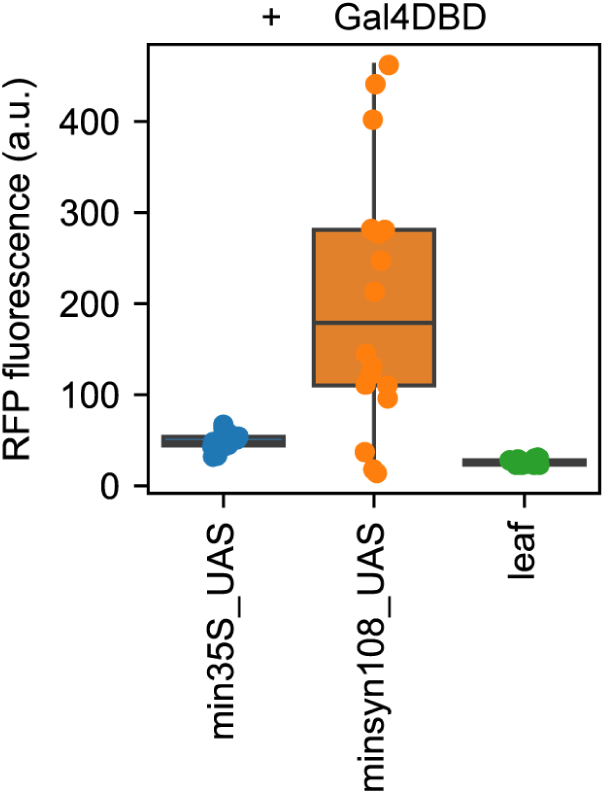
Basal expression level of different minimal promoters. Box plot of tissue-level RFP fluorescence of leaves infiltrated with Gal4DBD and either one of two minimal promoters driving RFP-T2A-SlWIP2-C NTF. The min35S_UAS promoter corresponds to the minimal -46 35S promoter containing 6 upstream Gal4 binding sites. The minsyn108_UAS promoter is a modified version of the previously published minsyn108 promoter containing 6 embedded Gal4 binding sites without deleting original sequences. Leaf autofluorescence is shown as a reference. The min35S_UAS promoter was used for all ENTRAP-seq experiments and for fluorescence reporter assays targeting activators. The minsyn108_UAS promoter was used for fluorescence assays to measure repressors in Fig.2F, Fig.3B. Boxes show median ± IQR, whiskers correspond to minimum and maximum values.

**Figure S5.**
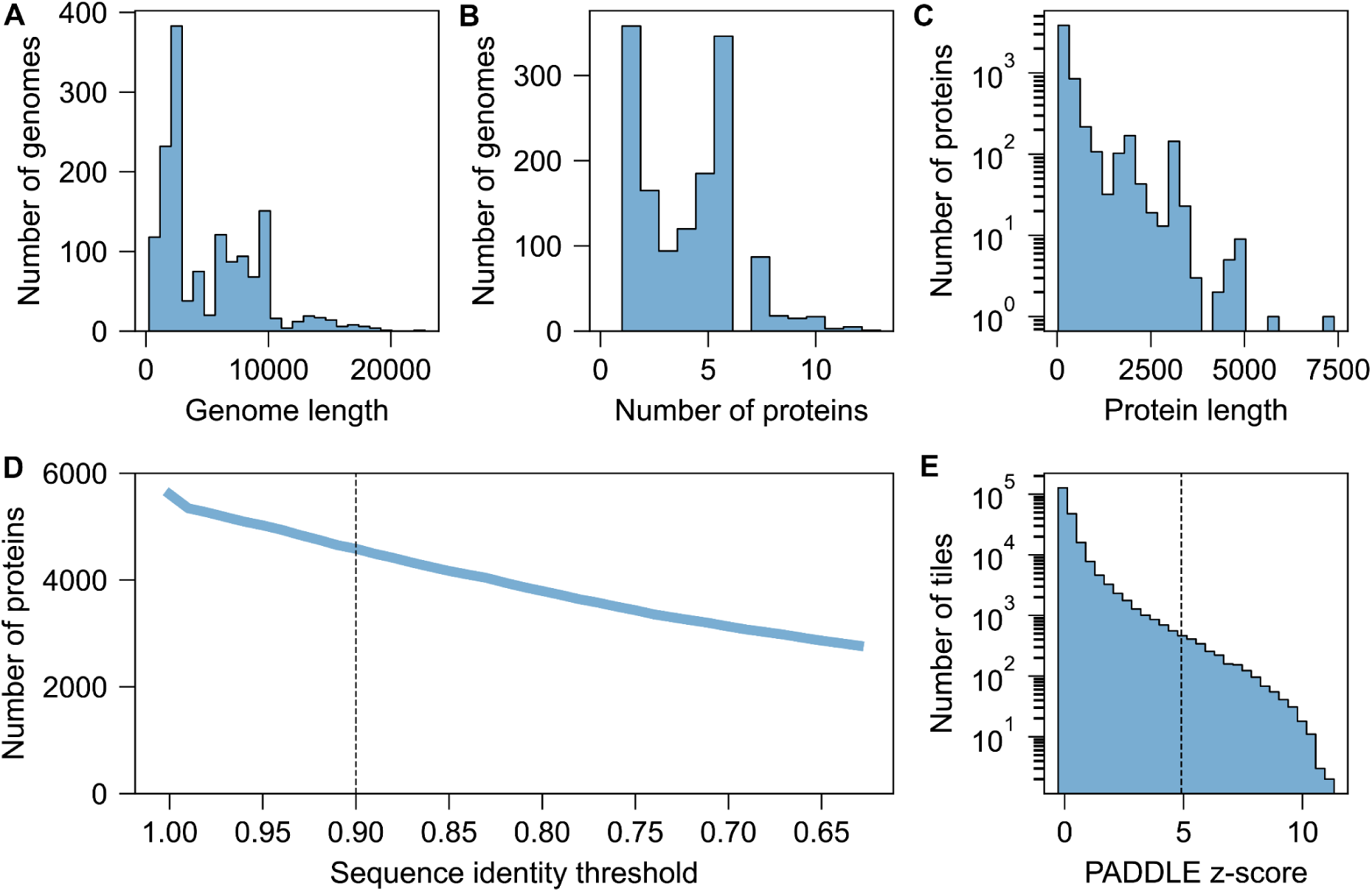
Overview of plant virus genomes and tile selection. Distribution of viral genome lengths. **(B)** Number of proteins in each viral genome. **(C)** Distribution of protein lengths. **(D)** Sequences were clustered at 90% identity (dashed line) to maximize protein diversity, and representative proteins were tiled for activator prediction using PADDLE. **(E)** The top 2,000 tiles with the highest PADDLE predicted z-score, corresponding to a threshold of 4.9 (dashed line), were selected for experimental characterization.

**Figure S6.**
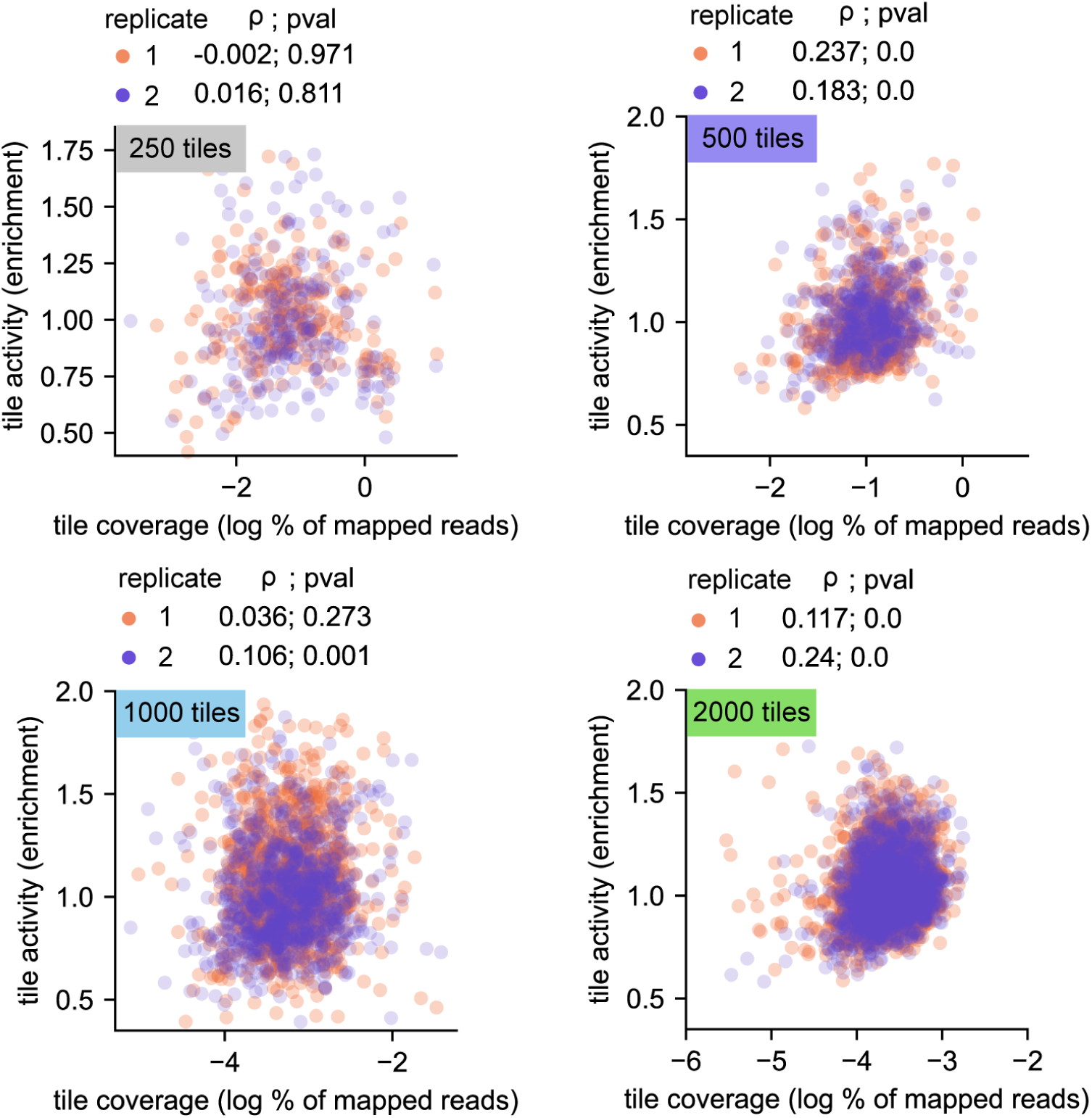
No correlation between tile coverage in the library and enrichment ratio. Scatter plots showing the transcriptional activity of each tile as measured by its enrichment value (y axis) as a function of the abundance of this tile in the library (x axis).

**Figure S7.**
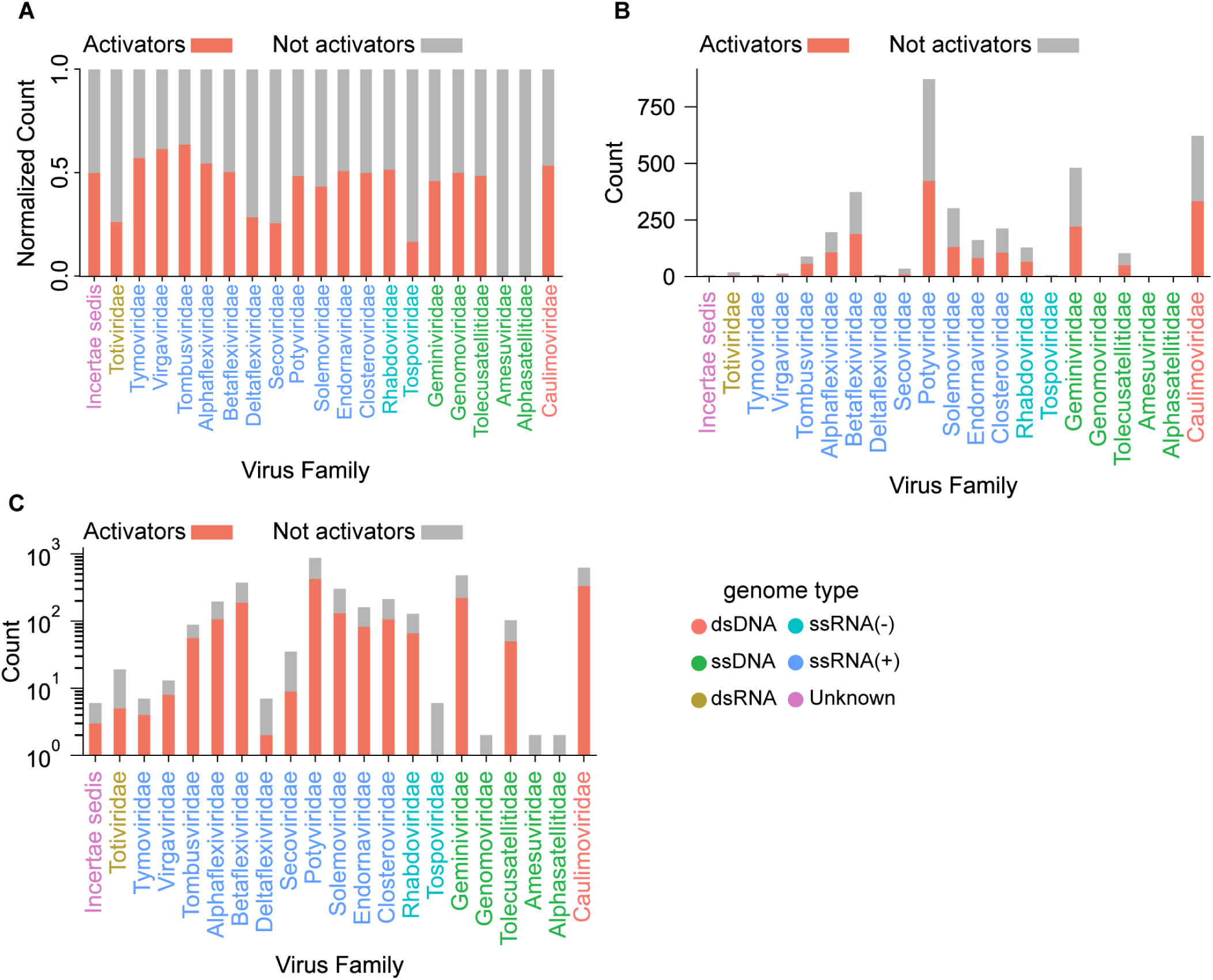
Fraction of selected tiles identified as activators across viral families. **(A)** For each virus family, the number of activator tiles is normalized by the total number of tiles selected for the corresponding family. **(B)** Same as (A) but showing absolute counts. **(C)** same as (B) showing the absolute counts in a log scale. Note that the viral families shown here do not exactly match the families in the tree of Fig. 4A. Some families such as *Totiviridae* had too few tested tiles to be displayed in the tree.Some families shown in the tree (e.g., *Amalgaviridae*) did not have any tiles that passed our criteria for testing.

**Figure S8.**
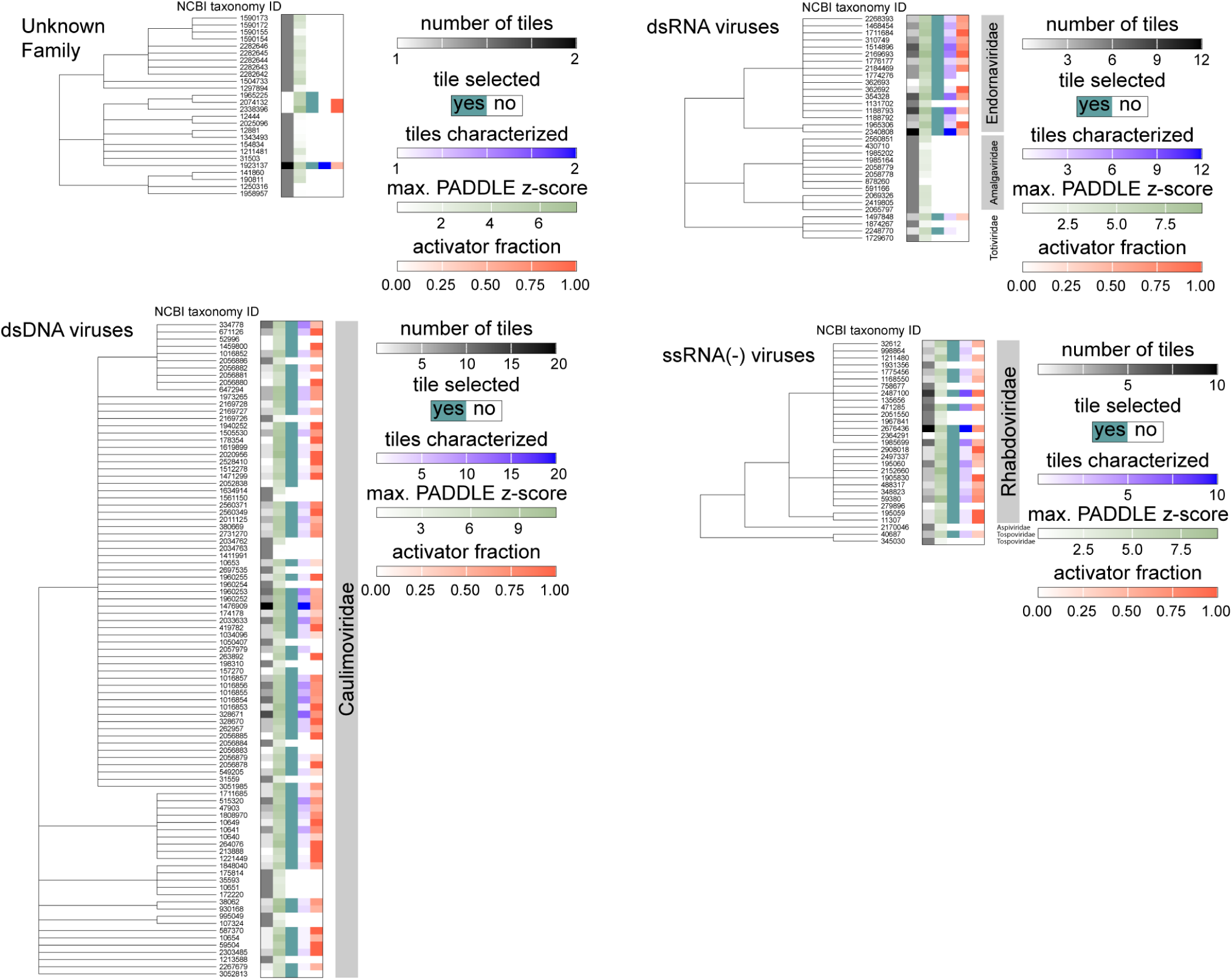
Details for double-stranded RNA, double-stranded DNA, single-tranded negative RNA, and unknown virus families. Shown are zoomed-in segments from the phylogenetic tree plot in Figure 4A. See also Table S1: Compiled data for viral libraries.

**Figure S9.**
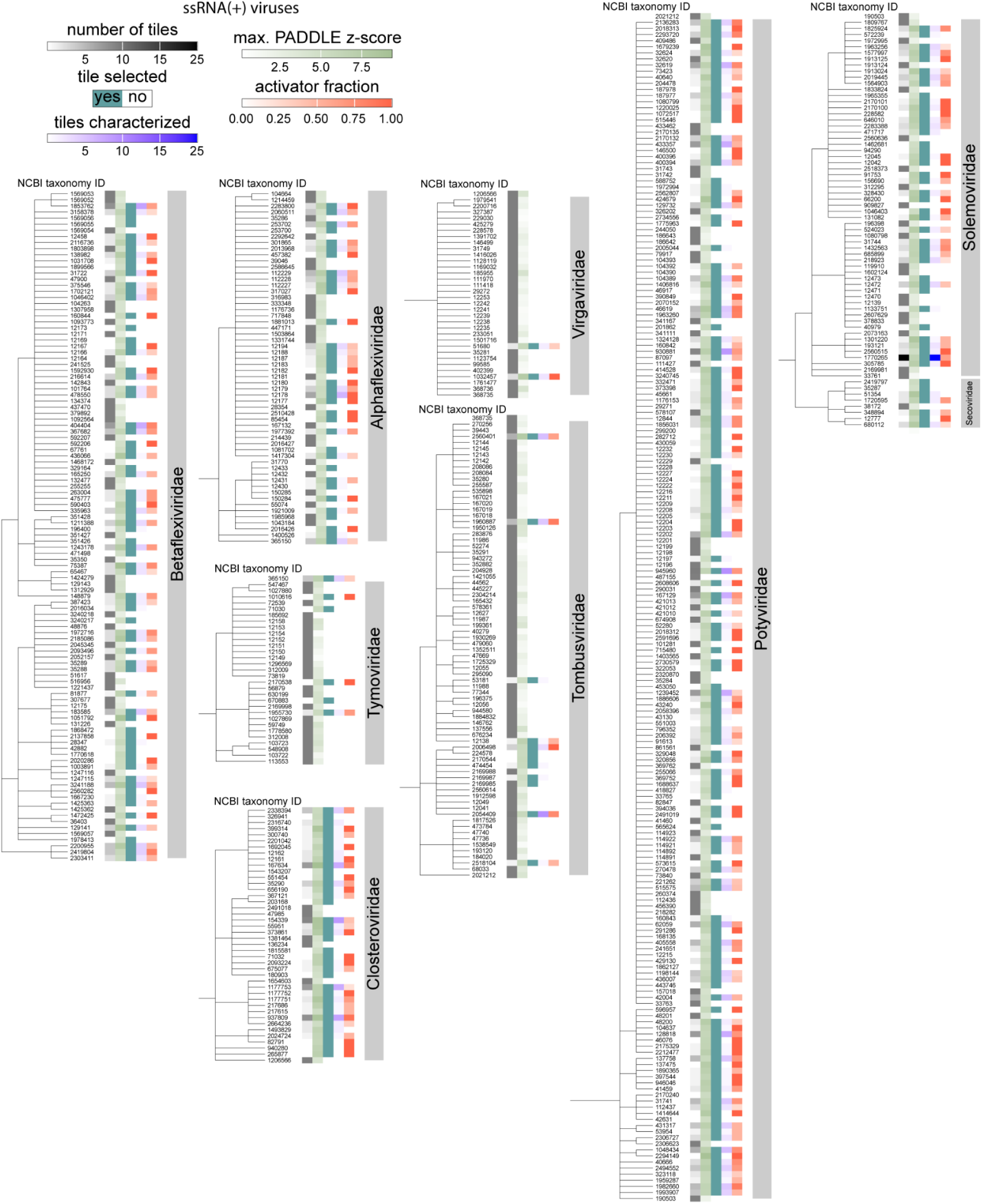
Details for positive single-stranded RNA virus families. Shown are zoomed-in segments from the phylogenetic tree plot in Figure 4A. See also Table S1: Compiled data for viral libraries.

**Figure S10.**
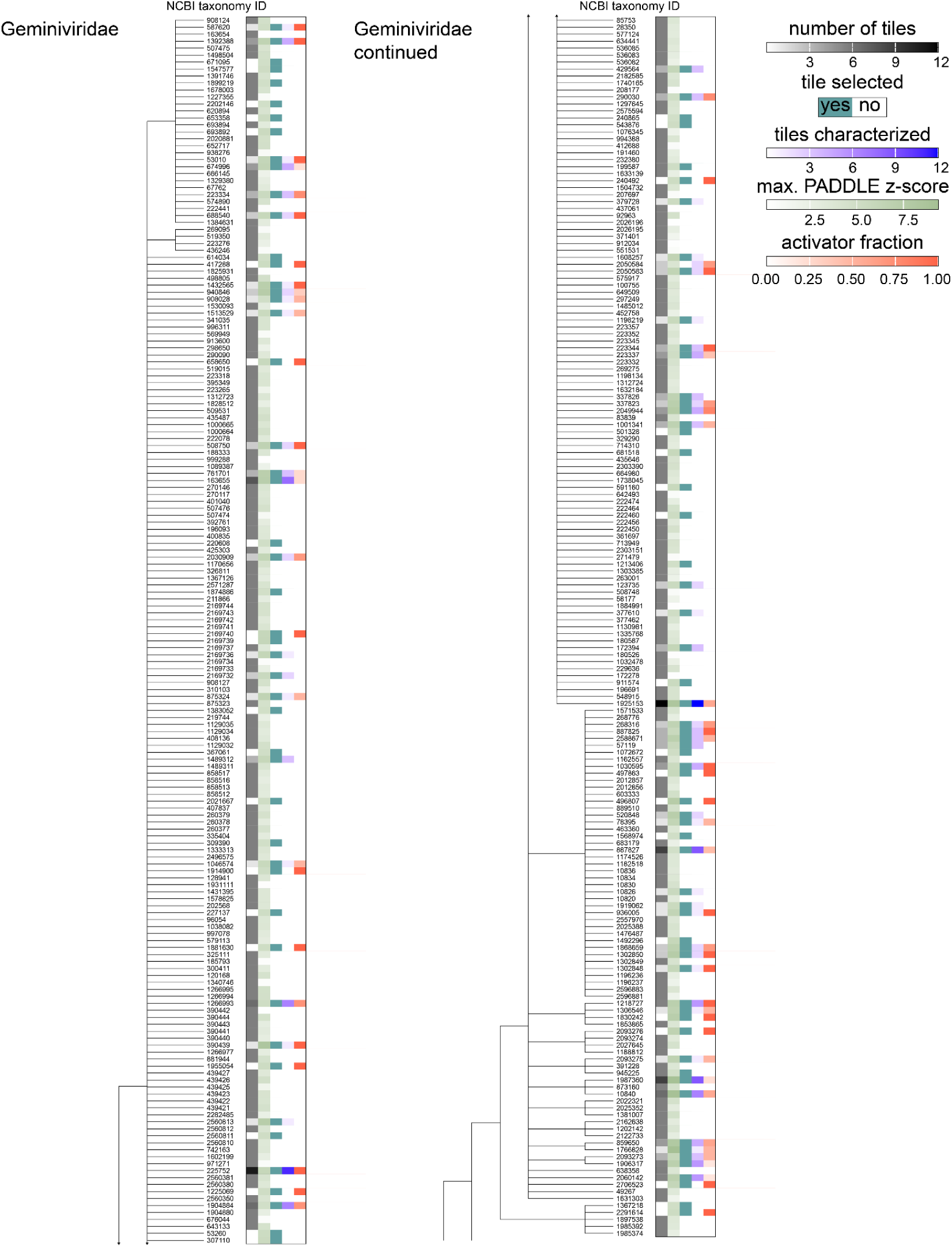
Details for *Geminiviridae*. Shown are zoomed-in segments from the phylogenetic tree plot in Figure 4A corresponding to viruses in the *Geminiviridae* family of single-stranded DNA viruses. See also Table S1: Compiled data for viral libraries.

**Figure S11.**
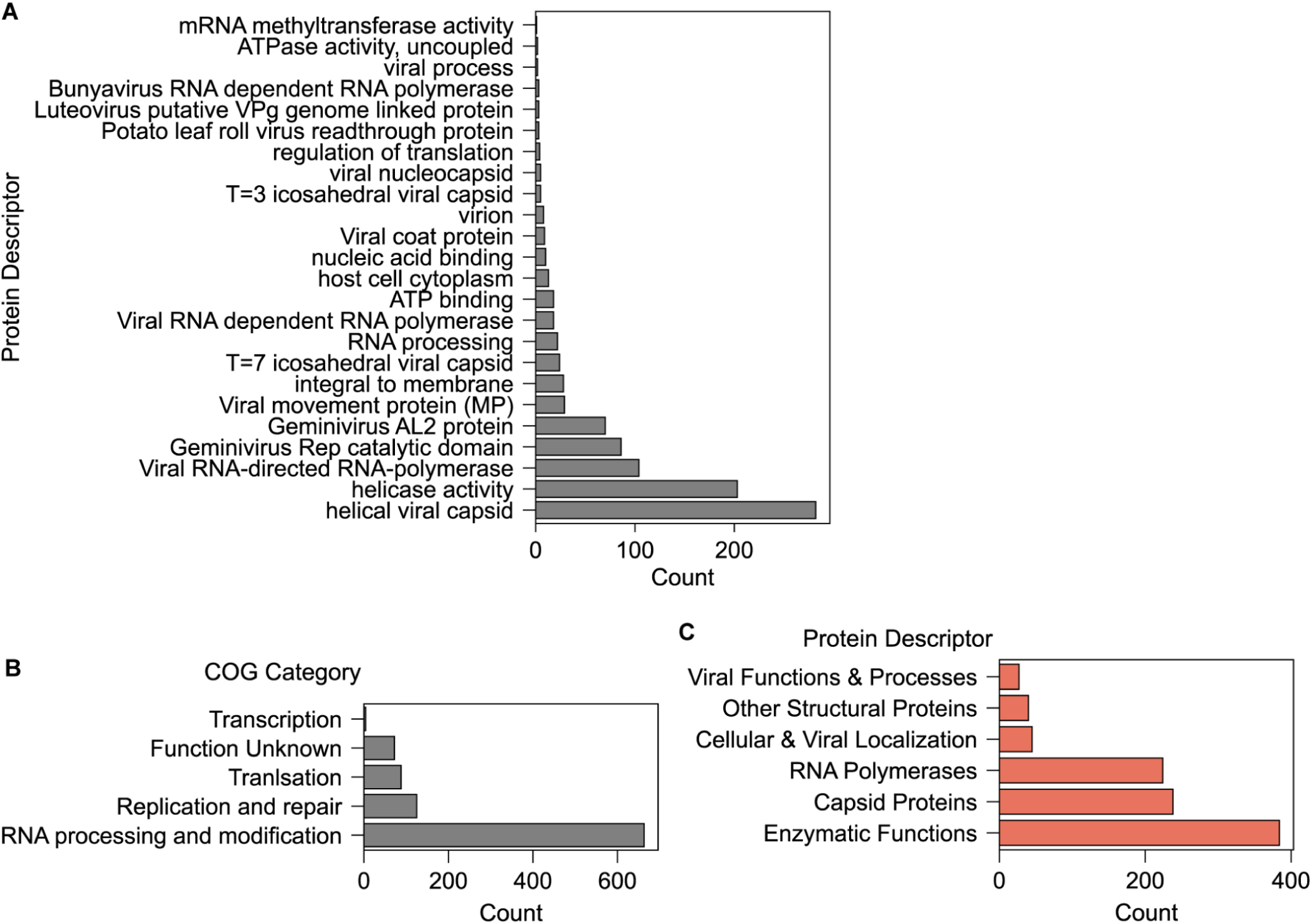
Functional annotation of proteins in the viral libraries. **(A)** GO term analysis across the whole library reveals enrichment of specific protein domains across library sequences, highlighting overrepresentation of functional classes among the tested viral protein sequences. **(B)** Clusters of Orthologous Groups (COG) categories in the whole library. **(C)** Protein descriptor categories among proteins for which we found an AD.

**Figure S12.**
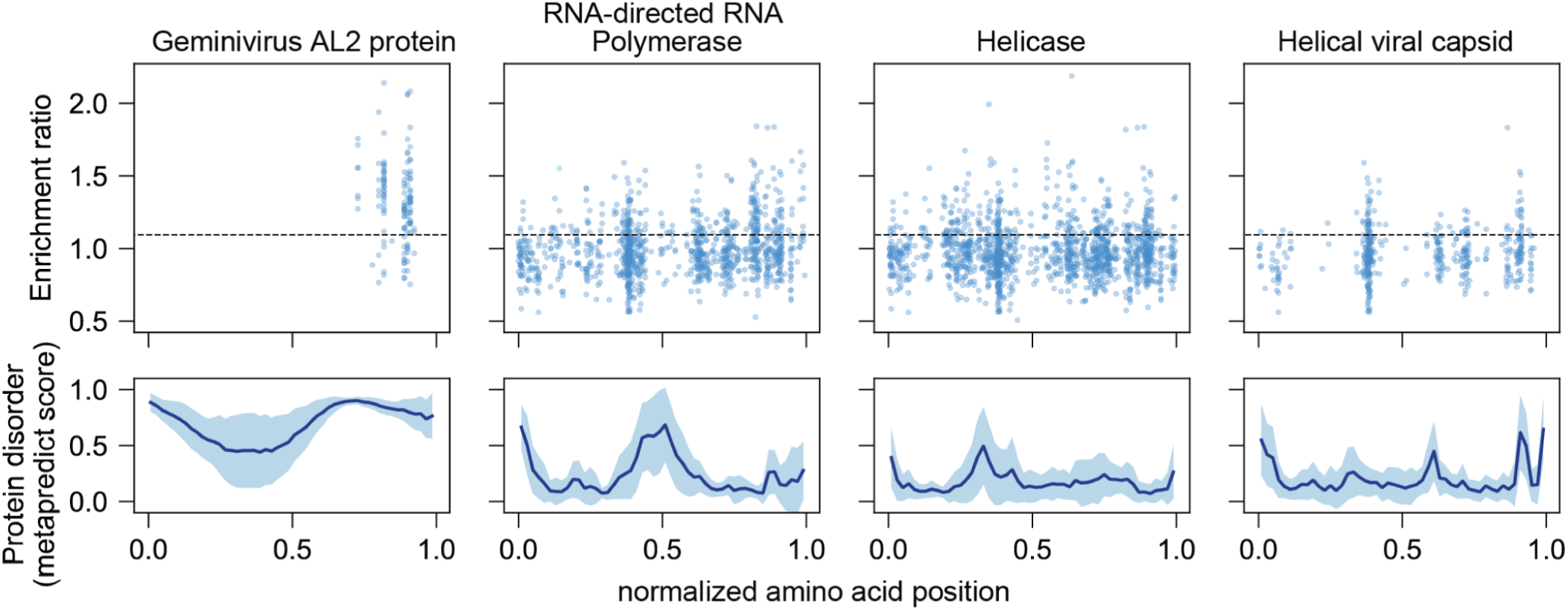
Enrichment ratios for tested tiles from four overrepresented protein classes. Top: Enrichment ratio of each tile along the (normalized) length of all proteins belonging to each given class. The amino acid position of each tile was normalized by dividing by the length of each corresponding protein. The horizontal dashed line shows the maximum threshold across datasets used to classify tiles as activators. Bottom: average ± 1 s.d. protein disorder as predicted by Metapredict.

**Figure S13.**
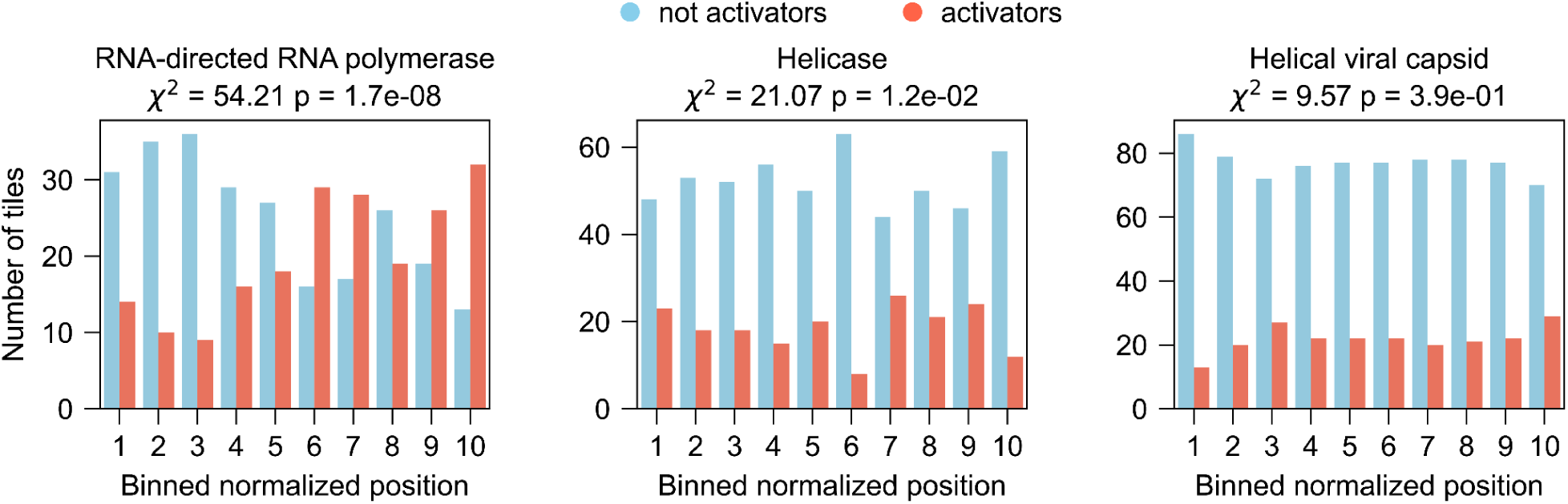
Positional clustering of putatively identified ADs. Normalized protein positions were binned and the distribution of identified AD regions showed statistically significant clustering in RdRp and helicase proteins, but not for helical capsid proteins, as determined by a chi-squared test.

**Figure S14.**
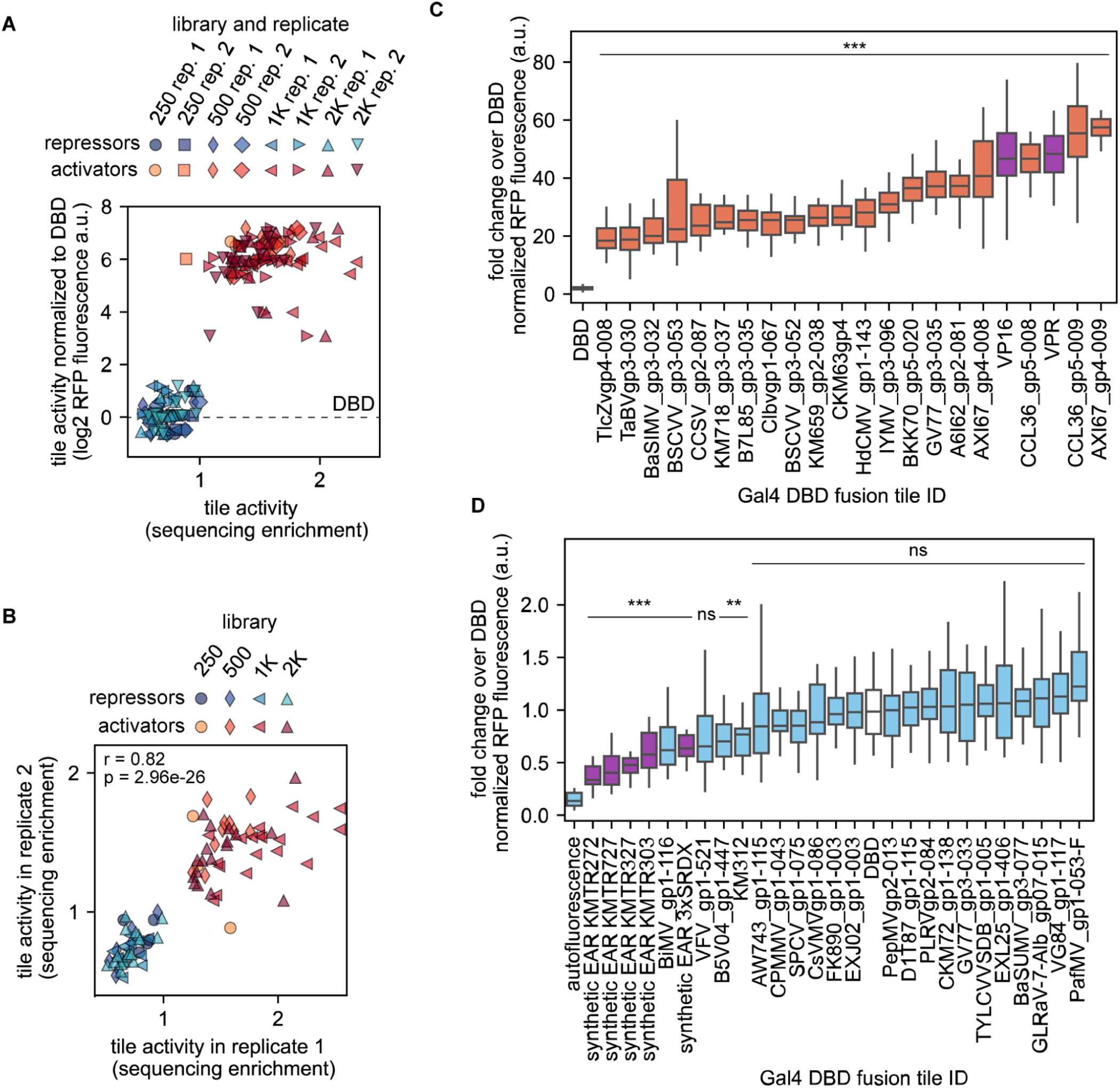
Testing potential strong activator and repressor tiles from plant viruses. **(A)** Data overview. 40 tiles with a high degree of enrichment (20 putative activators) or de-enrichment (20 putative repressors) in at least one library were selected for individual testing in a fluorescence reporter assay. Shown is the mean reporter fluorescence of each tile infiltrated individually (y-axis) as a function of tile enrichment for each replicate and library where this tile was present (x-axis). **(B)** Reproducibility of the data in (A). Scatter plot shows the enrichment of each candidate tile in two biological replicates. **(C)** Reporter fluorescence for each candidate activator tile expressed as a function of the DBD alone. Gal4 DBD fusions to VP16 and VP4 were included for comparison. **(D)** Reporter fluorescence for each candidate repressor tile expressed as a function of the DBD alone. Gal4 DBD fusions to synthetic EAR motifs were included for comparison. In (C) and (D), boxplots show the median and the interquartile range, whiskers show minimum and maximum values. *** p< 0.001, ** p<0.01, ns not significant of two-sided Welch’s t-tests against DBD with Bonferroni multiple comparison correction.

**Figure S15.**
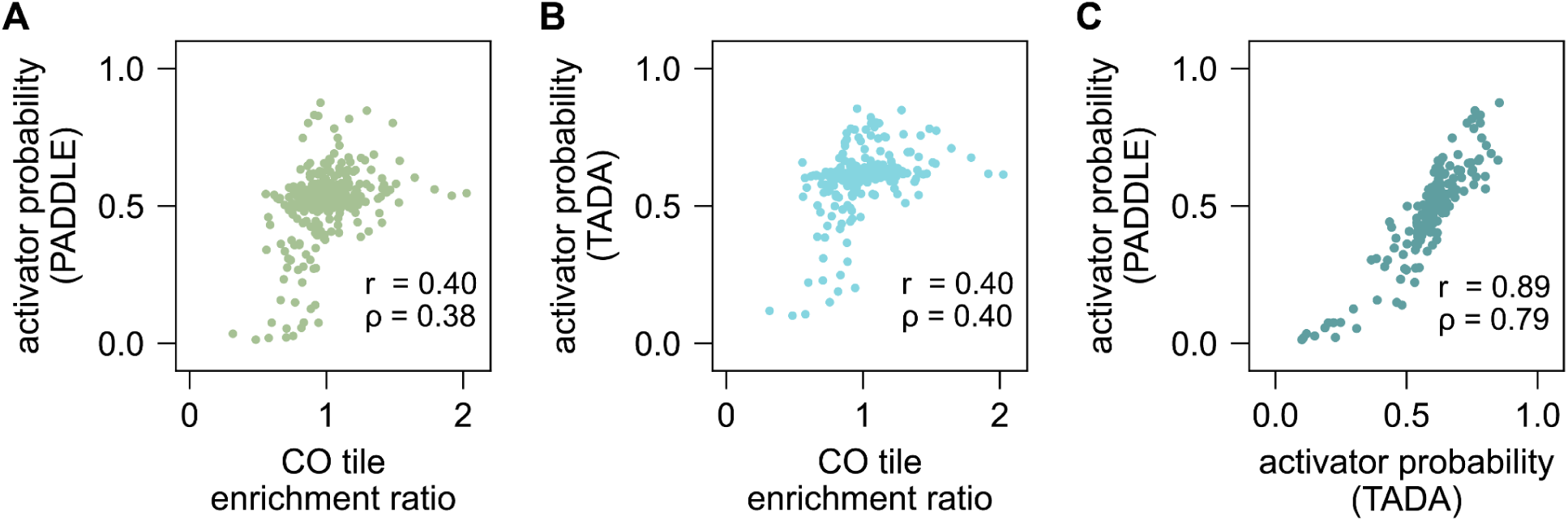
CO variant inference using PADDLE and TADA. **(A)** Median activator predictions from PADDLE and **(B)** TADA for each CO tile correlates with enrichment ratio. **(C)** Median PADDLE and TADA predictions for each CO tile also show strong concordance with respect to one another.

**Figure S16.**
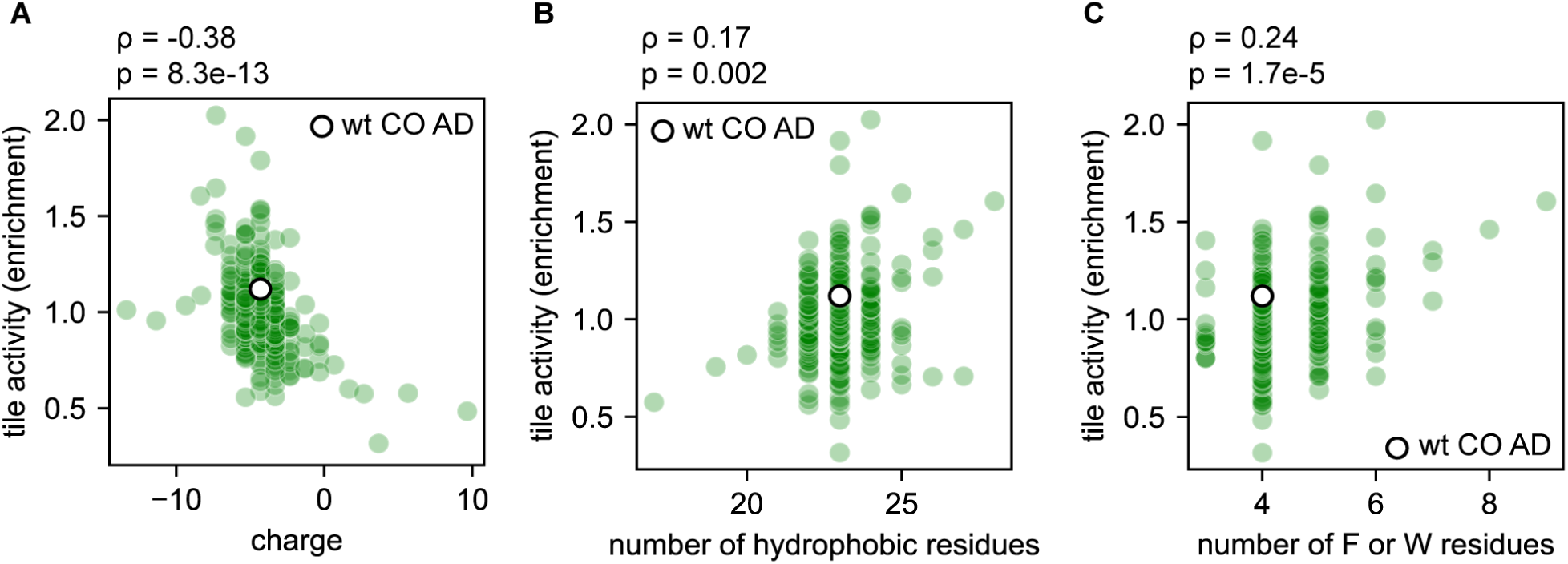
Biochemical properties of the CO acidic AD. The transcriptional activity of each tile in the CO AD library (y axis) is plotted as a function of different biochemical and sequence features. The wt tile is shown as a white circle. **(A)** Tile charge calculated as isoelectric point at pH 7.0. **(B)** number of hydrophobic residues. (C) Number of selected hydrophobic residues, phenylalanine (F) and tryptophan (W) and ⍴ = Spearman correlation coefficient.

**Figure S17.**
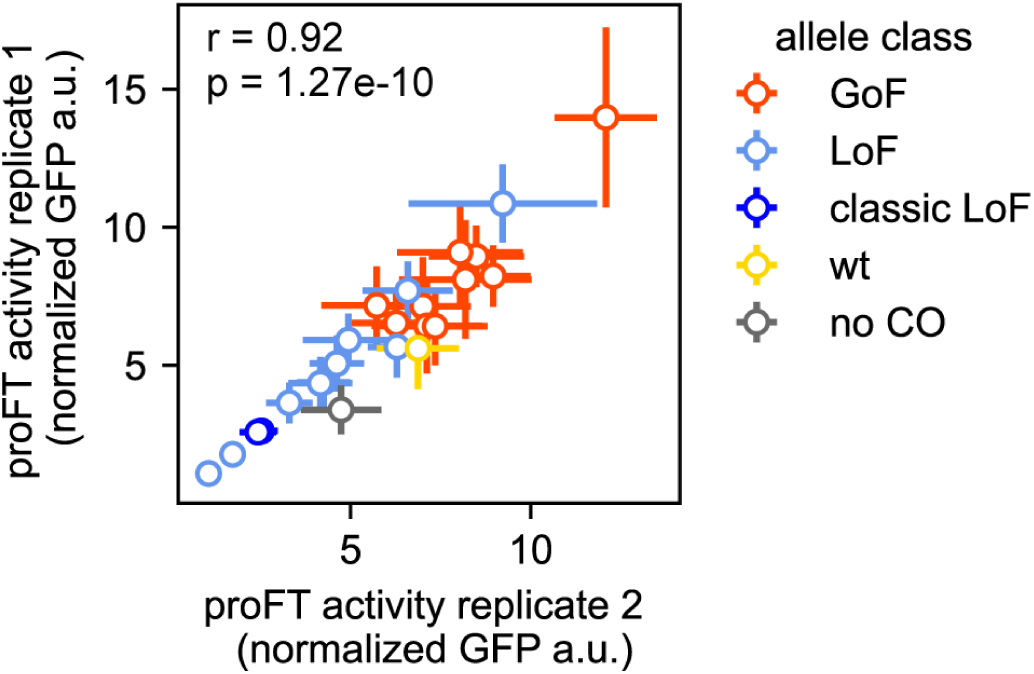
Replicate of CO alleles regulating the FT promoter driving GFP. The experiment shown in Fig. 6H was replicated on a different week. The FT promoter activity driven by a given CO allele on the first replicate (y axis) is plotted against the results from the same allele from the second replicate (x axis).

**Figure S18.**
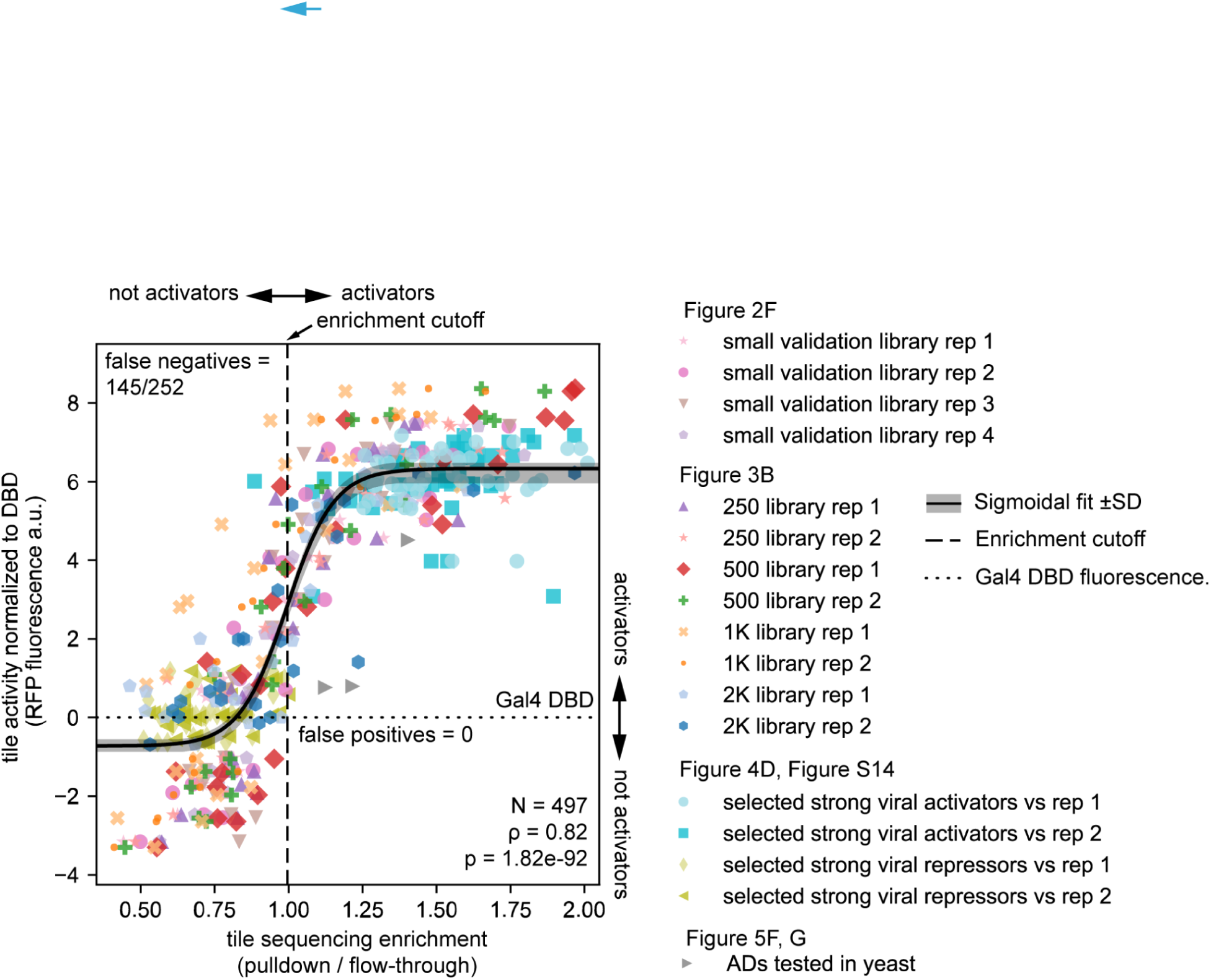
ENTRAP-seq quantitatively recapitulates fluorescence plate reader measurements. Scatter plot showing all presented cross validation experiments combined. Tile transcriptional activity based on individual infiltrations followed by fluorescence plate reader assays (y-axis) is compared against tile activity based on ENTRAP-seq experiments (enrichment, x-axis). Legend indicates where the data was originally presented in the text.

**Figure S19.**
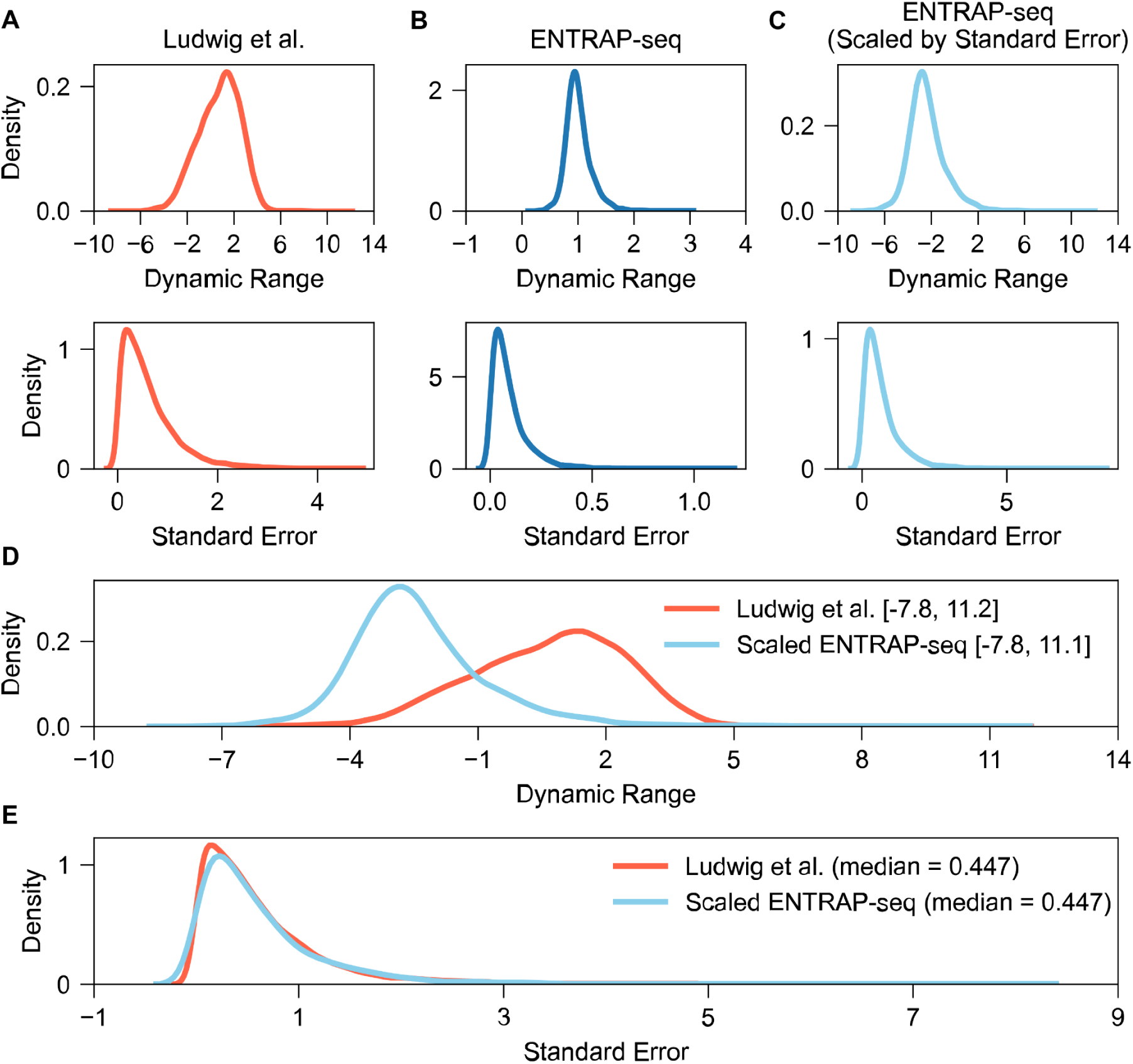
Measurement comparison between Ludwig et al. and ENTRAP-seq. **(A)** The dynamic range of measurements and their associated standard error for Ludwig et al ^22^. and **(B)** ENTRAP-seq. **(C)** Scaling the standard error of measurements in ENTRAP-seq to match the median standard error in Ludwig et al. results in a larger dynamic range with comparable distribution of standard error. **(D)** Scaling the dynamic range and shifting so that the minimum values are the same shows that the two technologies have comparable limits of detection and **(E)** standard error per measurement.

**Figure S20.**
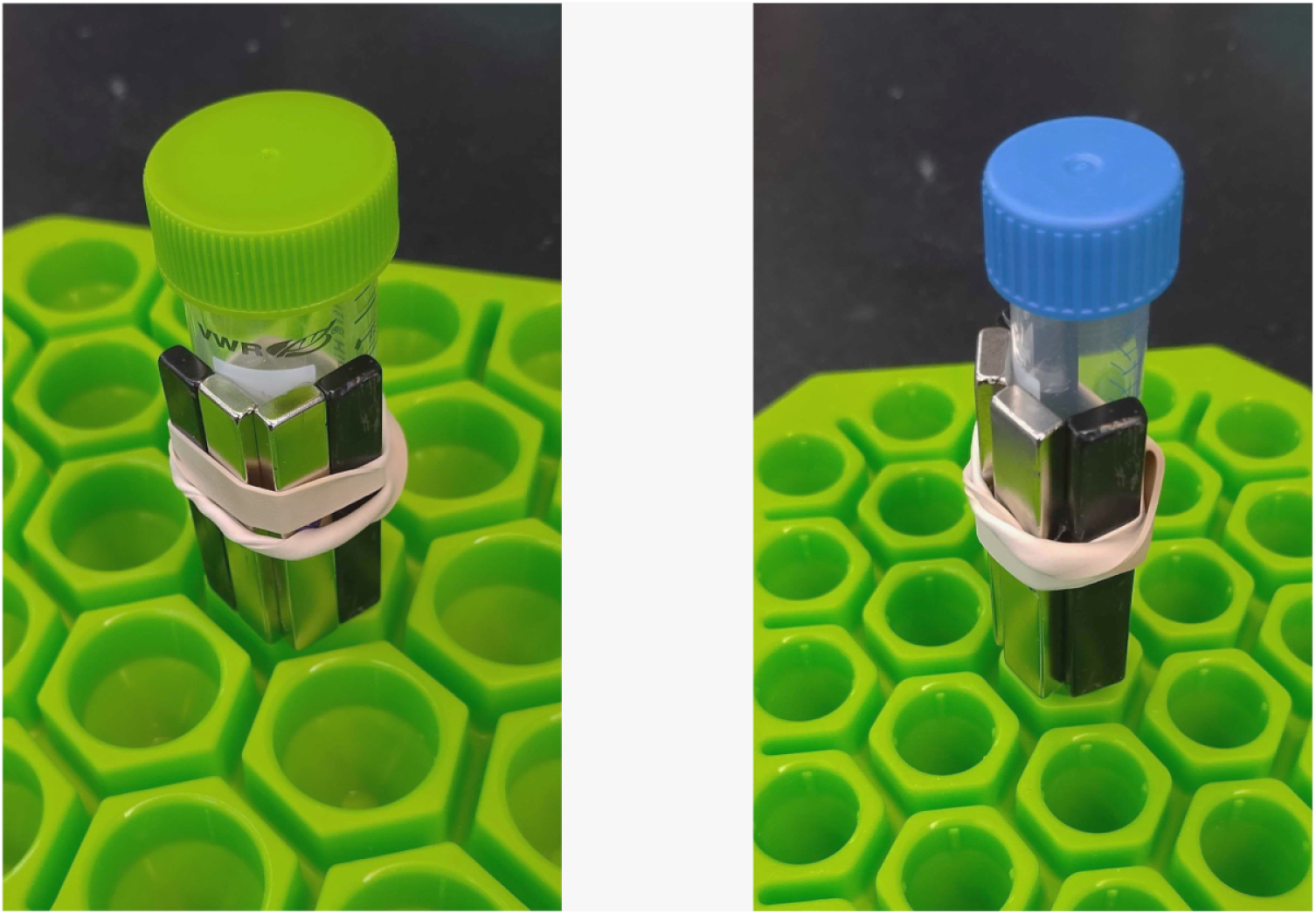
Setup for magnetic separation. Four neodymium magnets of dimensions 60 mm x 10 mm x 5 mm (LOVIMAG # B0CBTSFFSB) are attached to 50 ml (left) or 15 ml (right) falcon tubes using an elastic band.

## Supplementary Tables

All supplementary tables can be accessed from Zenodo (https://zenodo.org/records/16386135).

**Table S1:** Compiled data for viral libraries.

**Table S2:** Compiled data for yeast library.

**Table S3:** Compiled data for CO library.

**Table S4:** List of plasmids.

**Table S5:** List of primers.

